# Senescence inhibits the chaperone response to thermal stress

**DOI:** 10.1101/2021.06.15.448532

**Authors:** Jack Llewellyn, Venkatesh Mallikarjun, Ellen Appleton, Maria Osipova, Hamish TJ Gilbert, Stephen M Richardson, Simon J Hubbard, Joe Swift

## Abstract

Cells respond to stress by synthesising chaperone proteins that correct protein misfolding to maintain function. However, protein homeostasis is lost in ageing, leading to aggregates characteristic of protein-folding diseases. Whilst much is known about how these diseases progress, discovering what causes protein-folding to deteriorate could be key to their prevention. Here, we examined primary human mesenchymal stem cells (hMSCs), cultured to a point of replicative senescence and subjected to heat shock, as an *in vitro* model of the ageing stress response. We found through proteomic analysis that the maintenance of homeostasis deteriorated in senescent cells. Time-resolved analysis of factors regulating heat shock protein 70 kDa (HSPA1A) revealed a lack of capacities for protein turnover and translation to be key factors in limiting the stress response during senescence. A kinetic model predicted a consequence of these reduced capacities to be the accumulation of misfolded protein, a hypothesis supported by evidence of systematic changes to protein fold state. These results thus further our understanding of the underlying mechanistic links between ageing and loss of protein homeostasis.

## INTRODUCTION

Proteins are molecular machines required for cell and tissue function, providing structure and performing vital transport, signalling and enzymatic roles. The proteome must be actively regulated to match functional demands and address challenges – a state of proteostasis – thus ensuring that proteins are correctly folded, present in the right locations at the appropriate concentrations, and with any necessary post-translational modifications (Balchin et al, 2016; Sala et al, 2017). Due to its importance, proteostasis is safeguarded by a coordinated ‘proteostasis network’ (PN) that executes several functions: molecular chaperones assist in the folding of newly-synthesised proteins and resolve misfolding events as part of the cellular stress response, while protein degradation machinery allows misfolded or surplus proteins to be removed or recycled. Dysregulation of the PN is a recognised consequence of ageing (Hipp et al, 2019; Kaushik & Cuervo, 2015). Aged cells have decreased chaperone and proteasomal activity, and consequently accumulate oxidatively damaged and misfolded proteins (Koga et al, 2011). The detrimental effects of protein misfolding are two-fold: firstly, that loss of a protein’s structure leads to loss of its function; and secondly, that misfolded proteins can form aggregates that are toxic to cells. Protein aggregation is characteristic of diseases such as Alzheimer’s, Parkinson’s and Huntington’s – all disorders where age is considered a major risk factor (Hipp et al, 2014; Labbadia & Morimoto, 2015). Progress in the development of drugs to address diseases such as Alzheimer’s has been slow (Huang et al, 2020), but a better understanding of the PN may inform new therapeutic strategies.

The PN in human cells contains on the order of two thousand component proteins (Klaips et al, 2018). Within this group, ∼300 chaperone and cochaperone proteins (referred to as the ‘chaperome’) are associated with protein folding and conformational maintenance (Brehme et al, 2014). Chaperone proteins function by guiding their unfolded client proteins through a free energy landscape – traversing partially-folded intermediate states at points of local energetic minima – to find the global energetic minimum associated with the native folded state. In mechanistic terms, this process requires that chaperones shield exposed hydrophobic regions of their partially-folded clients, thus preventing unwanted interactions (Hartl et al, 2011). Many proteins within the chaperome are classified as heat shock proteins (HSPs) as they are expressed in response to heat as a prototypic form of stress, and are grouped into families by their molecular weight (kDa). Proteins within the HSP60, HSP70, HSP90 and HSP100 families have activity dependent on adenosine triphosphate (ATP) metabolism, whereas the small HSPs (sHSPs) are ATP-independent (Jayaraj et al, 2020). Chaperone function is further regulated by cochaperone proteins, such as the tetratricopeptide repeat (TPR) proteins that assist the HSP90 system and the HSP40 family of proteins that increase the client-specificity of HSP70 chaperones (Kampinga & Craig, 2010). The expression of many HSPs is coordinated by the heat shock response (HSR) pathway: here, an accumulation of misfolded protein causes disassembly of complexes of HSPs, allowing them to engage with client proteins. This disassembly process also releases the transcription factor heat shock factor 1 (HSF1), allowing it to translocate to the nucleus where it acts as a master regulator of HSP transcription.

More than one hundred genes related to the PN have been shown to be significantly suppressed with age in humans, including representatives of the chaperome from HSP40, HSP70 and HSP90 families (Brehme et al, 2014). Correspondingly, chaperone-assisted protein folding and disaggregation processes have been demonstrated to deteriorate with ageing (Calderwood et al, 2009; Koga et al, 2011). The protective response to heat stress was found to deteriorate with ageing in *C. elegans*, concomitant with increased levels of protein unfolding (Ben-Zvi et al, 2009). In the same model, knockdown of HSF1 was found to amplify age-associated loss of protein function, while its overexpression extended the maintenance of proteostasis and lengthened lifespan (Ben-Zvi et al, 2009; Hsu et al, 2003). In post-mitotic tissues of mice (heart, spleen, renal and cerebral cortices), basal levels of HSP70 protein were found to be decreased in aged vs. adult animals. Interestingly, however, adult levels of HSP70 were found to be maintained in naturally long-lived animals (de Toda et al, 2016). Expression of HSP70 has also been shown to increase in mice during exercise, a response thought to protect against muscle damage; overexpression of HSP70 reduced age-associated deterioration of muscle function (McArdle et al, 2003). A study of human-derived lymphoblasts has shown that the ability to increase levels of HSP70 gene expression in response to stress was generally decreased by ageing, but that the response was maintained in cells from exceptionally long-lived subjects (Ambra et al, 2004). Taken together, this evidence suggests that dysregulation of PN components with ageing is widely conserved, that age may affect both basal levels of HSPs and their ability to respond to stress, and that prolonged maintenance of proteostasis machinery may benefit both health and longevity. Nonetheless, a systematic survey of how the complex PN responds to stress, and how the response is affected by ageing, remains lacking – particularly in human cells and tissues.

Human ageing varies greatly between individuals due to genetic and environmental diversity, and therefore a lack of longitudinal ageing studies has limited our understanding of the process. Cellular senescence, such as achieved through replication of primary cells, has therefore often been used as an *in vitro* ageing model (Martinez Guimera et al, 2017). Senescence is a cellular response to irreparable DNA damage: to prevent propagation of potentially oncogenic damage the cell cycle is arrested and the cell marked for clearance. Changes to relative rates of cell damage, repair, clearance and renewal mean that aged tissues have disproportionately high numbers of senescent cells compared to young tissues (de Magalhaes & Passos, 2018; Lopez-Otin et al, 2013). Senescent cells have been reported to have decreased levels of HSPs (Deschenes-Simard et al, 2014), and lowered proteasomal (Saez & Vilchez, 2014) and mitochondrial activity (Martinez Guimera et al, 2017).

In this study, we have examined the response of primary human mesenchymal stem cells (hMSCs) to heat stress. This cell type has been widely studied in the context of tissue engineering and regenerative medicine, with a range of applications currently subject to clinical trials (Pittenger et al, 2019). This study has important implications to this purpose because: (i) how ageing and/or serial expansion in culture impacts on hMSC behaviour is an important consideration when applying autologous treatment strategies to older patients; and, (ii) many medical applications of hMSCs will require that the cells be robust to stress (Richardson et al, 2016), and inflammation, infection and tissue injury/repair are known triggers of HSF1-mediated stress response (Morimoto, 2008). We have used replicative senescence to model the effects of ageing, comparing early passage and senescent hMSCs from matched donors to maximise the statistical power of our analysis. By examining a combination of proteins and transcripts, we have constructed a systematic, time-resolved characterisation of how senescence affects the speed, magnitude and efficacy of the stress response. Furthermore, we have applied network and computational analysis to better understand the complexity of the PN and model the response to stress. We report on the senescence-induced decline of a functional module within the chaperome, centred around the activity of heat shock protein 70 kDa (HSPA1A), and attribute a weakening of the responsiveness to stress to depreciation of translational and protein turnover capacity. In summary, this investigation into the nature and cause of the decline of the HSR in senescent hMSCs provides mechanistic insight into how proteostasis is lost, and offers a starting point for strategies that could recover it.

## RESULTS

### The proteome is broadly affected by the onset of senescence

We first sought to characterise the intracellular proteomes of proliferating, early passage (EP) and senescent, late passage (LP) primary human mesenchymal stem cells (hMSCs) under control conditions (culture at 37 °C). EP hMSCs were used between passages 1 and 7, while onset of senescence in LP hMSCs occurred between passages 5 and 18 (varying between donors, see Table 1). Senescence was confirmed in LP cells by positive β-galactosidase staining (Supplementary Fig. S1A), expression of β-galactosidase protein (GLB1; *p* < 0.0001, false discovery rate (FDR)-corrected ANOVA; Supplementary Fig. S1B), and loss of lamin B1 protein (LMNB1; *p* < 0.0001, FDR-corrected ANOVA) and transcript (*LMNB1; p* < 0.0001, t-test; Supplementary Figs. S1B, C) (Shimi et al, 2011). LP cells were also found to have greater spread areas than EP cells, characteristic of senescence (*p* < 0.0001, ANOVA; Supplementary Fig. S1D) (Hernandez-Segura et al, 2018). Log_2_ fold-changes in protein level were quantified in whole cell lysates using liquid chromatography-coupled tandem mass spectrometry (LC-MS/MS) (Mallikarjun et al, 2020), comparing donor-matched LP vs. EP hMSCs; of 1830 proteins identified with ≥3 peptides-per-protein, 286 were found to be significantly upregulated, and 520 significantly downregulated (*p* < 0.05, FDR-corrected ANOVA; Fig. 1A). Ontological analysis using the Reactome database (Fabregat et al, 2017; Fabregat et al, 2018; Jassal et al, 2020) showed significant downregulation of “cell cycle”, “DNA replication” and “translation” pathways (FDR-corrected *p*-values < 0.05), suggesting suppression of processes consistent with the ‘hallmarks of cellular senescence’ (Hernandez-Segura et al, 2018). Interestingly, both “cellular response to heat stress” and “regulation of HSF1-mediated heat shock response” pathways were significantly diminished in LP vs. EP hMSCs, even in the absence of heat stress.

**Table 1.** Information on primary human mesenchymal stem cell (hMSC) donors.

| Donor ID | Donor gender | Source site | Donor age | Passage number<br>(proliferating, “EP”) | Passage number<br>(senescent, “LP”) |
| --- | --- | --- | --- | --- | --- |
| WH143 | M | Hip | 58 | 2 | 7 |
| WH176 | F | Hip | 64 | 2 | 7 |
| TH191 | F | Hip | 61 | 4 | 9 |
| TH194 | M | Knee | 65 | 3 | 8 |
| WH211 | F | Hip | 36 | 7 | 18 |
| TH215 | M | Knee | 73 | 3 | 5 |
| TH223 | M | Knee | 64 | 4 | 7 |
| WH226 | F | Hip | 54 | 2 | 9 |
| WH230 | M | Hip | 43 | 3 | 7 |
| TH269 | M | Knee | 70 | 1 | 7 |
| TH296 | M | Hip | 68 | 2 | 6 |
| TH2305 | M | Knee | 67 | 3 | 7 |
| TH309 | F | Knee | 69 | 4 | 10 |

**Figure 1.**
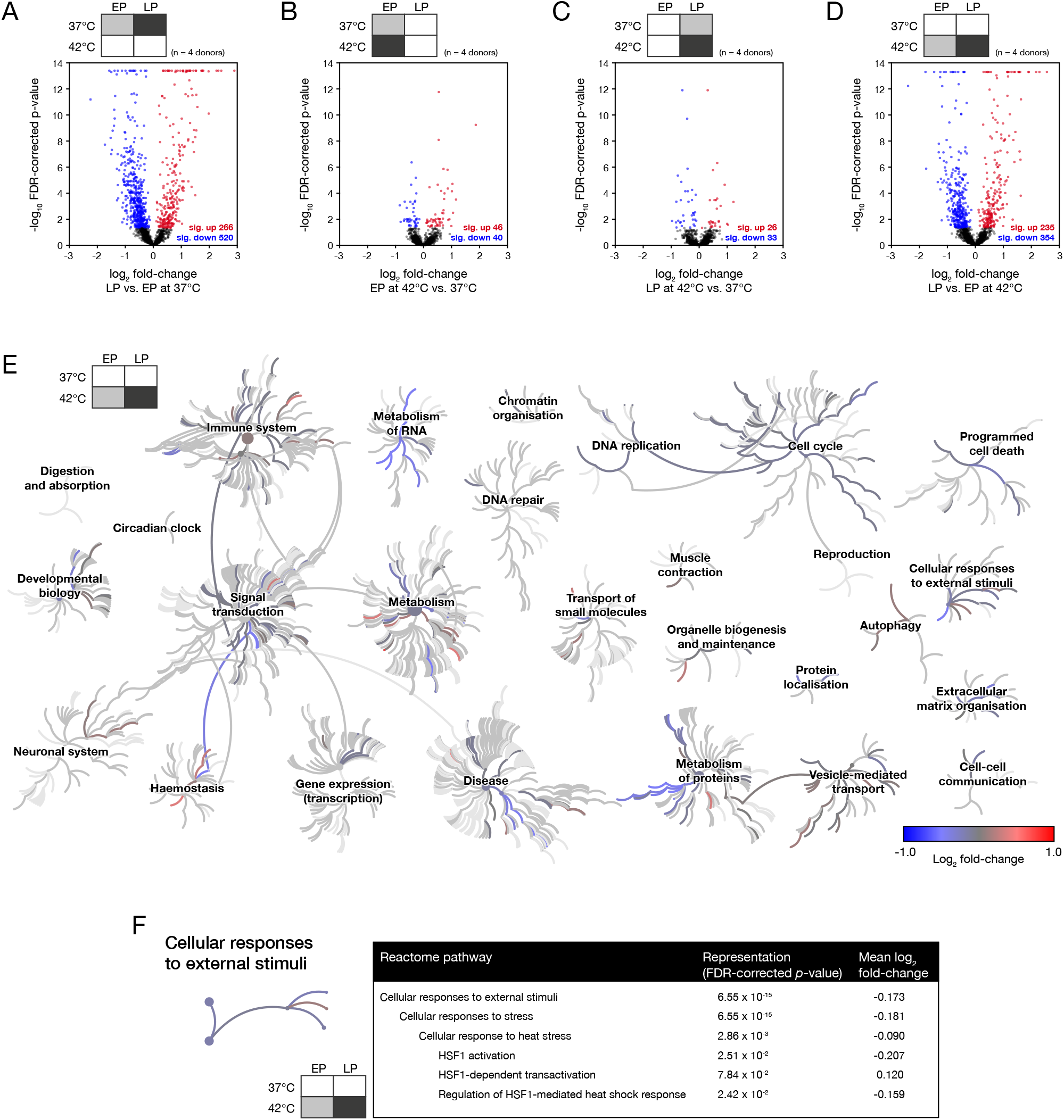
Proteomic analysis shows that senescent human mesenchymal stem cells (hMSCs) have a supressed response to thermal stress. Mass spectrometry was used to compare the proteomes of donor-matched early passage (EP) and senescent late passage (LP) hMSCs with and without a 2-hour heat shock treatment at 42 °C; whole-cell lysates were then analysed using mass spectrometry proteomics. Volcano plots showing the distribution of changes in the abundance of 1830 proteins in (**A**) LP vs. EP hMSCs in the absence of heat shock; (**B**) EP hMSCs with and without heat shock; (**C**) LP hMSCs with and without heat shock; (**D**) LP vs. EP hMSCs, both subjected to heat shock. In figure panels (A)-(D), red and blue points satisfy a *p*-value < 0.05. *p*-values were calculated using empirical Bayes-modified t-tests with Benjamini–Hochberg false discovery rate (FDR) correction (Mallikarjun et al, 2020); *n* = 4 primary donors. (**E**) Reactome pathway analysis of the significantly affected proteins shown in panel (D), comparing the responses of EP and LP hMSCs subjected to heat shock (Fabregat et al, 2017; Fabregat et al, 2018; Jassal et al, 2020). Significantly represented pathways (false discovery corrected *p*-value < 0.05) are shown in colours corresponding to the mean log_2_ fold-change of proteins in the pathway. (**F**) Expanded view of the pathway “Cellular response to heat stress” and its parent and child pathways. The “Cellular response to heat stress” pathway was found to be significantly supressed in the response of LP cells to heat shock, with a false discovery rate (FDR) corrected *p*-value of 2.86 × 10^−3^.

### The response to heat stress is attenuated in senescent cells

Having characterised the effects of senescence on the hMSC proteome, we examined the responses of EP and LP cells to heat stress treatment. Using LC-MS/MS to again quantify changes to 1830 proteins detected with ≥3 peptides-per-protein, we compared hMSCs subjected to a 2-hour treatment at 42 °C to matched cells under control conditions. Using EP hMSCs, we found that heat stress caused significant upregulation of 46 proteins and downregulation of 40 proteins (*p* < 0.05, FDR-corrected ANOVA; Fig. 1B). The number of significantly-affected proteins was decreased in LP hMSCs subjected to the same treatment: 26 upregulated and 33 downregulated (*p* < 0.05, FDR-corrected ANOVA; Fig. 1C). Examination of affected Reactome pathways showed that “cellular response to heat stress” and “regulation of HSF1-mediated heat shock response” were significantly upregulated in response to heat stress in both EP and LP hMSCs (FDR-corrected *p*-values < 0.05), however, the mean fold-change to protein levels in these pathways was decreased in LP cells, indicative of a suppressed stress response. Notably, the perturbation to the proteome caused by heat shock to EP or LP hMSCs was considerably smaller than the change following the onset of senescence (comparing Figs. 1A-C). We therefore compared the proteomic endpoints of both EP and LP hMSCs following the heat treatment, as means of contrasting their relative capacities to manage proteotoxic stress. In addition to generating a volcano plot of LP vs. EP hMSCs following 2-hour heat treatment at 42 °C (Fig. 1D), we mapped the mean log_2_ fold-changes of proteins within significantly affected Reactome pathways onto a network capturing all cellular processes (FDR-corrected *p*-values < 0.05, Fig. 1E). This plot again highlights broad proteomic differences between EP and LP hMSCs – attributed to processes such as metabolism, cell cycle, apoptosis and organisation of the extracellular matrix – but also suggests a failure to adequately respond to proteotoxic stress following senescence: the parent pathway “cellular responses to external stimuli” was significantly affected (FDR-corrected *p*-values < 0.0001; shown expanded in Fig. 1F), and within that, both “cellular response to heat stress” and “regulation of HSF1-mediated heat shock response” were suppressed.

### The human chaperome can be divided into functional modules based on protein-protein interactions

In order to better understand the consequences of suppressed stress response in senescent cells, we sought to characterise the roles of individual functional modules within the chaperome. The human chaperome described by Brehme et al. consists of 332 proteins; inclusion within this network of chaperone and cochaperone proteins was based on analysis of the literature, domain structures and annotations of human and *C. elegans* genomes (Brehme et al, 2014). Using the highest-confidence protein-protein interaction (PPI) data taken from the STRING database (Szklarczyk et al, 2019), we performed modularity analysis (Newman, 2006) on the Brehme et al. chaperome to subdivide it into deeply interconnected modules (Supplementary Figs. S2A, B). The question we sought to answer was – in physical terms – whether all chaperones had the broad remit of maintaining proteostasis as a collective (i.e. low modularity), or whether chaperones could be separated into functional modules that perform more specialised tasks (i.e. high modularity). We found the human chaperone network to be highly modular, dividing into 19 communities (Fig. 2A). We next analysed functional annotations within the individual interacting communities, focusing on the five modules most represented by our LC-MS/MS data. These functional modules did indeed seem to be dedicated to specific tasks within the maintenance of proteostasis. For instance, nearly all members of the HSP40 and HSP70 families of proteins were grouped together into one module (“HSP70 machinery”), while chaperones localised in the endoplasmic reticulum (ER) were grouped together in a module containing members of the P4HA and PLOD families of proteins (“ER-specific chaperones”).

**Figure 2.**
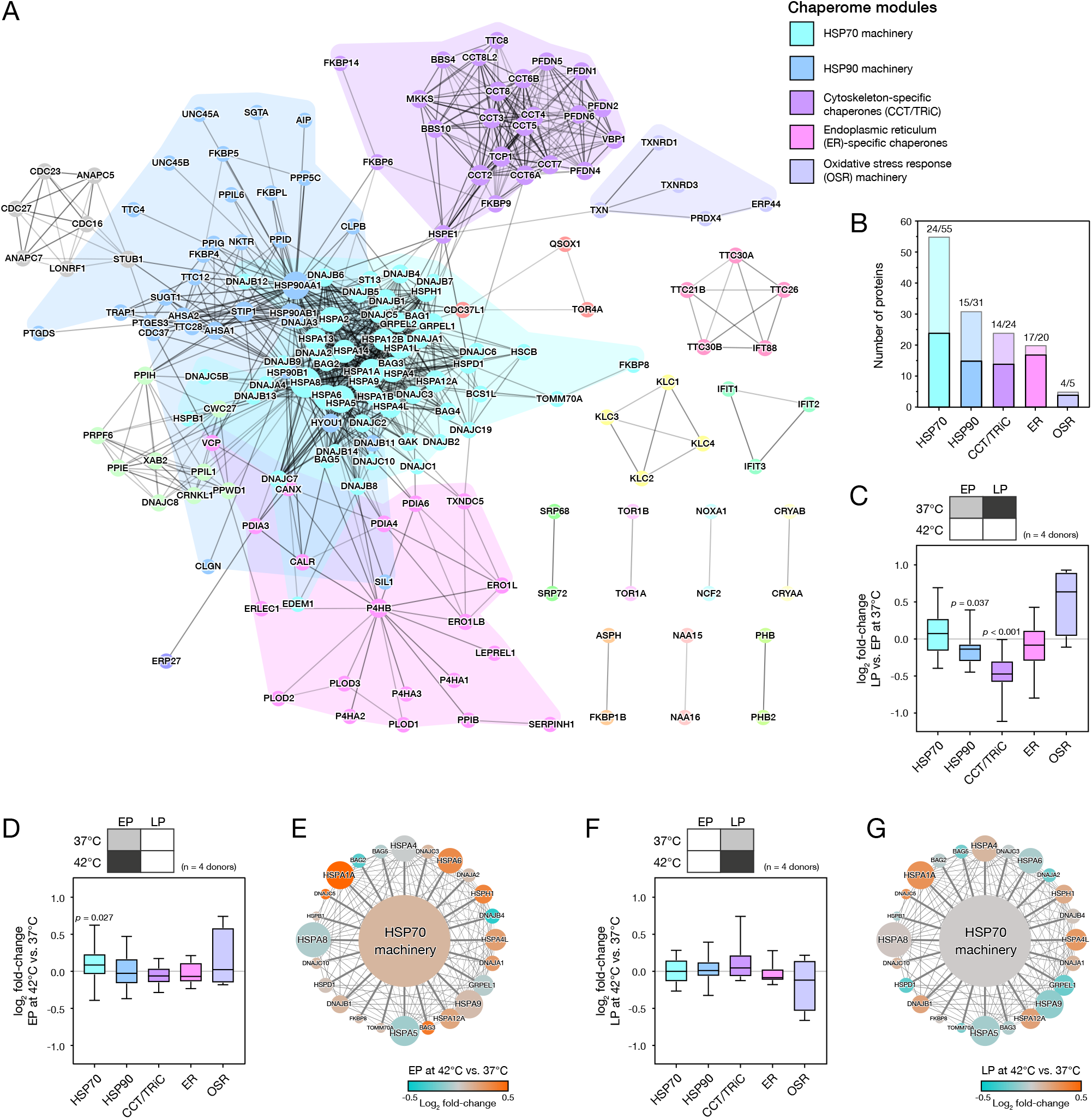
Network analysis of the chaperome shows that the heat-stress response of a heat shock protein 70 kDa (HSP70) module is lost in late passage (LP) human mesenchymal stem cells (hMSCs). (**A**) The community structure of proteins identified in the human chaperone network (Brehme et al, 2014). Proteins (nodes) are coloured by their inclusion into modules, with the five largest modules named according to their function: heat shock protein 70 kDa (HSP70) machinery (55 proteins); heat shock protein 90 kDa (HSP90) machinery (31 proteins); cytoskeleton-specific proteins (chaperonin containing t-complex / TCP1 ring complex, CCT/TRiC; 24 proteins); endoplasmic reticulum (ER)-specific chaperones (20 proteins); and, oxidative stress response (OSR) machinery (5 proteins). Node size is indicative of chaperone degree (Supplementary Fig. S2A), while edge weight indicates the STRING interaction score between chaperones (Supplementary Fig. S2B) (Szklarczyk et al, 2019). (**B**) Plot showing the number of constitutive proteins from each chaperome module detected in the mass spectrometry dataset presented in Fig. 1. (**C**) Fold-changes to proteins in chaperome modules in donor-matched LP vs. EP hMSCs in the absence of heat shock treatment. (**D**) Fold-changes to proteins in chaperome modules in EP hMSCs subjected to a 2-hour heat shock treatment at 42 °C. The HSP70 module was significantly upregulated (*p* = 0.027). (**E**) Network showing changes to individual proteins in the HSP70 module of figure panel (D). (**F**) Fold-changes to proteins in chaperome modules in LP hMSCs subjected to heat shock; in contrast to the EP cells, none of the modules were significantly regulated. (**G**) Changes to individual proteins in the HSP70 module of figure panel (F). In figure panels (C), (D) and (F), box-whisker plots show medians, quartiles and range; *p*-values were determined by ANOVA testing, *n* = 4 primary donors. In panels (E) and (G), node size is indicative of chaperone degree, while edge weight indicates interaction score between chaperones. The central node is coloured according to the mean abundance change of chaperones associated with the HSP70 machinery.

### A module based around HSP70 machinery becomes unresponsive to heat stress in senescent cells

Having established how the chaperome could be subdivided into interacting functional groups, we reanalysed our initial proteomic characterisation of the HSR in EP and LP hMSCs (Fig. 1). We found that the 1830 proteins detected with ≥3 peptides-per-protein in our proteomics dataset gave representative coverage of the five main chaperome groups identified by modularity analysis (minimum coverage of 43%, Fig. 2B). Analysis of the data comparing LP vs. EP hMSCs in the absence of heat stress showed significant downregulation of two functional chaperome modules in senescence: HSP90 machinery (*p* = 0.037, ANOVA), and cytoskeleton-specific proteins (chaperonin containing t-complex / TCP1 ring complex, CCT/TRiC; *p* < 0.001, ANOVA) (Fig. 2C). Given the role of proteins constitutive of the CCT/TRiC module in maintenance of the cytoskeleton, its substantial downregulation in senescence may be of consequence to the mechanical changes that occur to cells and tissues during ageing (Phillip et al, 2015). Nonetheless, when we examined the response of EP hMSCs to heat stress, we found only the module associated with HSP70 machinery to have been upregulated (*p* = 0.03, ANOVA; Fig. 2D). The HSP70 machinery module contains proteins from the HSPA family that were found to have the highest number of interactions in our modularity analysis, indicating that they are the key nodes within the chaperome network (Supplementary Fig. S2A) (Rubinov & Sporns, 2010). Heat shock cognate 71 kDa protein (HSPA8) had 64 protein-protein interactions, the greatest number in the network; heat shock protein 70 kDa (HSPA1A) had 47 network interactions, and was also the most stress-sensitive protein within the module (Fig. 2E). In contrast, when senescent hMSCs were subjected to the same heat stress treatment, there was no significant response in any of the chaperome modules (Fig. 2F); in comparison to EP cells, fold changes to proteins constitutive of the HSP70 machinery module were generally supressed (Fig. 2G). Together, this evidence demonstrates that in EP hMSCs, upregulation of HSP70 machinery is the central feature in cellular management of the proteotoxic shock caused by heat stress, but that senescent hMSCs are no longer capable of this response.

### Early passage cells are robust to inhibition of chaperone activity in the absence of thermal stress

We next examined regulation of the proteome in cells subjected to targeted inhibition of chaperone machinery. The small molecule 2-phenylethynesulfonamide (PES) has been shown to bind selectively to HSPA1A, inhibiting its activity by preventing interactions with its cochaperones (Leu et al, 2017; Leu et al, 2009; Leu et al, 2011). We found that treatment of EP hMSCs with PES did not greatly perturb the proteome: of 1830 proteins detected with ≥3 peptides-per-protein, only 26 proteins were significantly upregulated, and 22 downregulated (*p* < 0.05, FDR-corrected ANOVA; Fig. 3A). The five major chaperome modules identified in Figure 2A were also not significantly affected (ANOVA, Fig. 3B), suggesting that the system was robust to PES treatment in the absence of additional stress factors. Nonetheless, we did identify proteins within the HSP70 module that were possibly upregulated as part of a compensatory mechanism, such as heat shock 70 kDa protein 6 (HSPA6) and ER chaperone BiP (HSPA5) (Fig. 3C).

**Figure 3.**
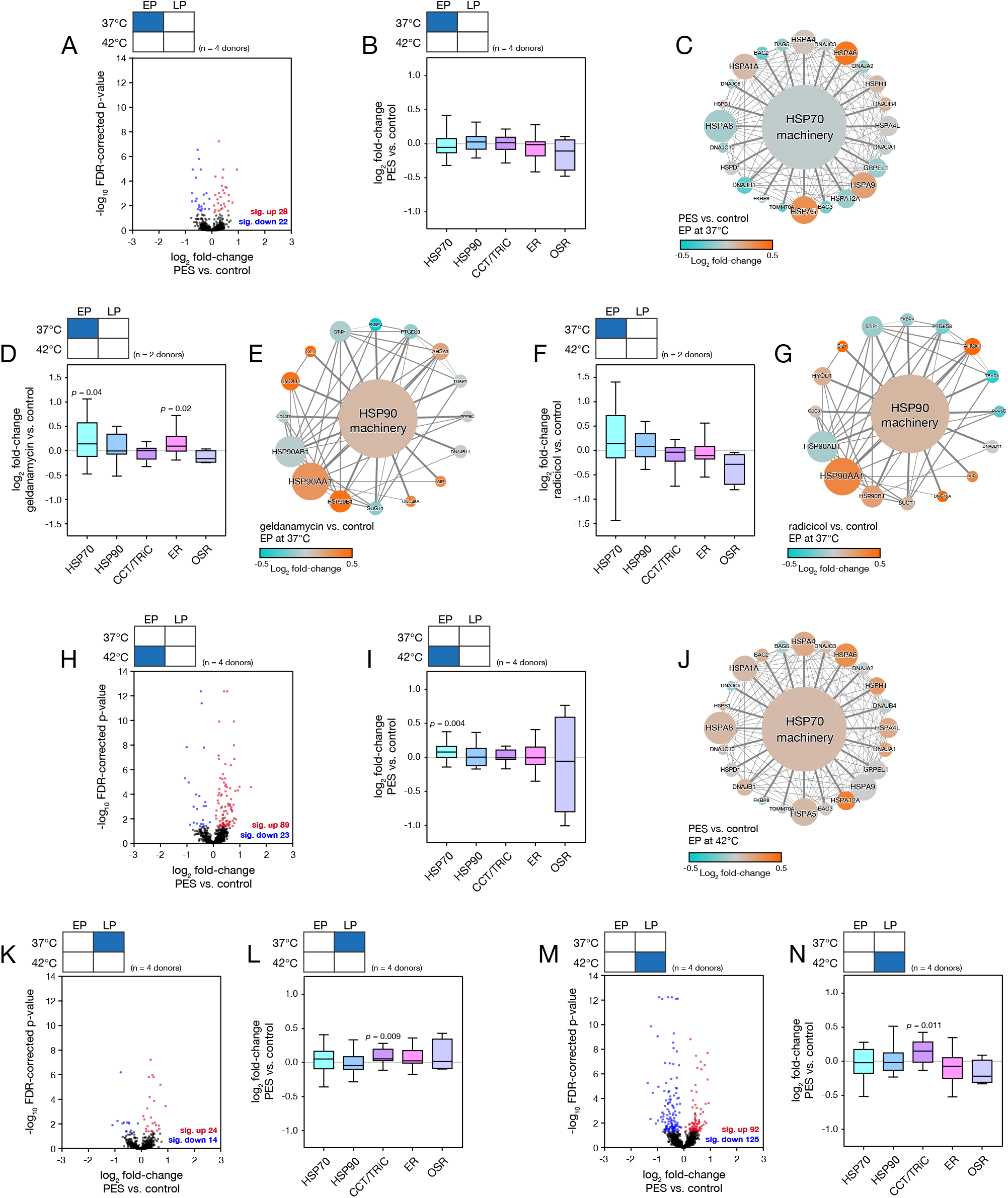
Early passage (EP) human mesenchymal stem cells (hMSCs) exhibit compensatory mechanisms when treated with chaperone inhibitors, but this response is dampened in late passage (LP) cells. (**A**) Volcano plot showing the effect of HSP70-inhibitor 2-phenylethynesulfonamide (PES) on EP hMSCs. (**B**) PES treatment caused no significant changes to chaperome modules in EP hMSCs. (**C**) Network showing changes to individual proteins in the HSP70 module of figure panel (B). (**D**) and (**E**) Perturbation to the chaperome and proteins in the HSP90 module caused by treatment with HSP90-inhibitor geldanamycin. (**F**) and (**G**) Perturbation to the chaperome and HSP90 caused by treatment with HSP90-inhibitor radicicol. (**H**) Volcano plot showing the effect of PES on EP hMSCs subjected to a 2-hour heat shock at 42 °C. (**I**) EP hMSCs treated with PES in combination with heat shock exhibited significant upregulation of the HSP70 module (*p* = 0.004). (**J**) Network showing changes to individual proteins in the HSP70 module of figure panel (I). (**K**) Volcano plot showing the effect of PES on LP hMSCs under control conditions. (**L**) Analysis of changes to chaperome modules in the data shown in (K); the HSP70 module was not significantly affected, but proteins in the cytoskeleton-specific chaperome module (CCT/TRiC) were significantly increased (*p* = 0.009). (**M**) Volcano plot showing the effect of PES on LP hMSCs subjected to 2-hour heat shock treatment at 42 °C. (**N**) Chaperome module analysis of data in (M); the HSP70 module was again unaffected, while the CCT/TRiC module were significantly increased (*p* = 0.011). In volcano plots, red and blue points satisfy a *p*-value < 0.05. *p*-values were calculated using empirical Bayes-modified t-tests with Benjamini–Hochberg false discovery rate (FDR) correction (Mallikarjun et al, 2020); *n* = 4 primary donors. Significance of changes to modules was determined by ANOVA testing. In network diagrams, node size is indicative of chaperone degree, while edge weight indicates interaction score between chaperones. The central node is coloured according to the mean abundance change of chaperones associated with the module.

To test the generality of this apparent robustness, we also examined the effects of targeted inhibition of HSP90 machinery. Geldanamycin and radicicol are both able to bind to the N-terminal nucleotide-binding domain of HSP90 proteins, preventing critical ATPase activity (Roe et al, 1999). Treatment of EP hMSCs with geldanamycin did not greatly perturb the proteome (of 1841 proteins detected with ≥3 peptides-per-protein, 10 were significantly upregulated and 22 downregulated; Supplementary Fig. S3A), but both HSP70 and ER chaperome modules were upregulated (*p* = 0.04 and *p* = 0.02, respectively, ANOVA; Figs. 3D, E). Treatment with radicicol had a broader effect on the proteome (30 significantly upregulated and 118 downregulated proteins; Supplementary Fig. S3A), but this response was less specific to the chaperome as no functional modules were significantly perturbed (ANOVA, Figs. 3F, G). In similarity with the response to HSP70 inhibition, neither of the HSP90 inhibitors caused a significant overall perturbation to their target chaperome module (i.e. HSP90 machinery), but we did observe common changes to individual proteins within the module, consistent with a compensatory response: both drug treatments resulted in increased levels of HSP90AA1 (the inducible form of HSP90; the constitutively expressed HSP90AB1 was not affected), HSP90 cochaperones AHSA and UNC45A, and ER-resident proteins endoplasmin (HSP90B1) and nucleotide exchange factor SIL1 (Figs. 3E, G).

### Senescence increases sensitivity to heat stress when combined with inhibition of HSP70

Having found EP hMSCs to be robust to inhibition of key chaperone proteins in the absence of stress, we applied the same proteomic tools to examine the effects of combining HSP70 inhibition with heat shock and senescence. Analysis of the effect of PES treatment on EP hMSCs subjected to a 2-hour treatment at 42 °C showed a broader perturbation to the proteome than PES-treated cells maintained at 37 °C (comparing Fig. 3H to Fig. 3A): 89 proteins were significantly upregulated, and 23 downregulated (*p* < 0.05, FDR-corrected ANOVA). The chaperome module related to HSP70 machinery was significantly upregulated (*p* = 0.004, ANOVA; Fig. 3I), and quantities of individual protein components within the module were generally increased (Fig. 3J). This result mirrors the response of EP hMSCs to heat stress in the absence of PES (Figs. 2D, E), interpretable as the cells responding to proteotoxic stress, finding the response ineffective due to the inhibition of HSP70, and therefore increasing its magnitude. A comparison between LP hMSCs with and without PES treatment showed a muted response. Only 24 proteins were significantly upregulated and 14 downregulated (*p* < 0.05, FDR-corrected ANOVA; Fig. 3K), and the HSP70 chaperome module was not significantly affected (Fig. 3L, Supplementary Fig. S3C). This is similar to the result found in EP cells (Figs. 3A-C), as in the absence of stress, inhibition of specific stress response machinery seemed to have a limited impact. In contrast, the effect of PES treatment on senescent hMSCs subjected to a 2-hour treatment at 42 °C was more substantial: 92 proteins were significantly upregulated, and 125 downregulated (*p* < 0.05, FDR-corrected ANOVA; Fig. 3M). However, this perturbation was not reflected in the HSP70 chaperome module, which was not significantly affected (Fig. 3N), and many of the individual proteins within the module appeared to be suppressed (Supplementary Fig. S3D). This is again reminiscent of earlier experiments performed in the absence of PES, where LP cells were unable to mount a chaperome response to thermal stress (Figs. 2F, G). Interestingly, PES treatments on LP cells at both temperatures caused an upregulation of the CCT/TriC chaperome module (Figs. 3L, N), notably suppressed in senescence (Fig. 2C). This hints at crosstalk between chaperome modules, and a possible role for HSP70 machinery in cytoskeletal maintenance. Taken together, these results highlight the failure of senescent cells to respond to proteotoxic stress in the same way as early-passage cells. Furthermore, this dysregulation was not caused by a desensitization to temperature – the proteome of LP cells was more widely affected by PES treatment at elevated temperature – but rather a failure to remodel specific features of the chaperome.

### Senescence suppresses the expression of HSPA1A protein following heat stress

In order to build a mechanistic understanding of the difference between the stress response in EP and LP hMSCs, we undertook to examine key components of the HSP70 machinery with finer temporal resolution. HSPA1A was visualised by immunofluorescence (IF) microscopy before, halfway through, and immediately following a 2-hour heat treatment at 42 °C, and at timepoints over a 24-hour recovery period (Fig. 4A). Subsequent quantification of mean HSPA1A IF intensity (analogous to protein concentration in the cell) showed that protein levels increased significantly during the heat treatment, reached a maximum 4 hours into the recovery period, and then slowly began to decrease (Fig. 4B). This was the case in both EP and LP hMSCs, with the levels of HSPA1A prior to heating not differing significantly. However, initial production of HSPA1A was delayed by approximately an hour in the LP response, and reached a significantly lower maximum – only 63% of that found in EP cells (see Figs. 4B, C for significance values, determined by ANOVA). An interesting property of HSPA1A is that it is known to be rapidly and reversibly translocated to the nucleus in cells subjected to thermal stress (Velazquez & Lindquist, 1984; Welch & Feramisco, 1984). This process, mediated by the transport protein HIKESHI (meaning ‘firefighter’), is thought to protect the contents of the nucleus from proteotoxic damage (Kose et al, 2012). By separately quantifying HSPA1A levels in the nucleus and cytoplasm (demarcated by DAPI and phalloidin staining, Fig. 4A), we found that nuclear HSPA1A levels mirrored those characterised in the rest of the cell: they reached a maximum after 4 hours, and levels were lower in LP than EP hMSCs (Supplementary Figs. S4A, B). We also examined the ratio between nuclear and cytoplasmic HSPA1A levels, finding significant differences between the behaviours of EP and LP cells, both during heat treatment and in the following 2 hours (Supplementary Figs. S4C, D). This is suggestive of changed patterns of client proteins, or regulation of transport processes, and consequently that senescence may affect how chaperone resources are directed.

**Figure 4.**
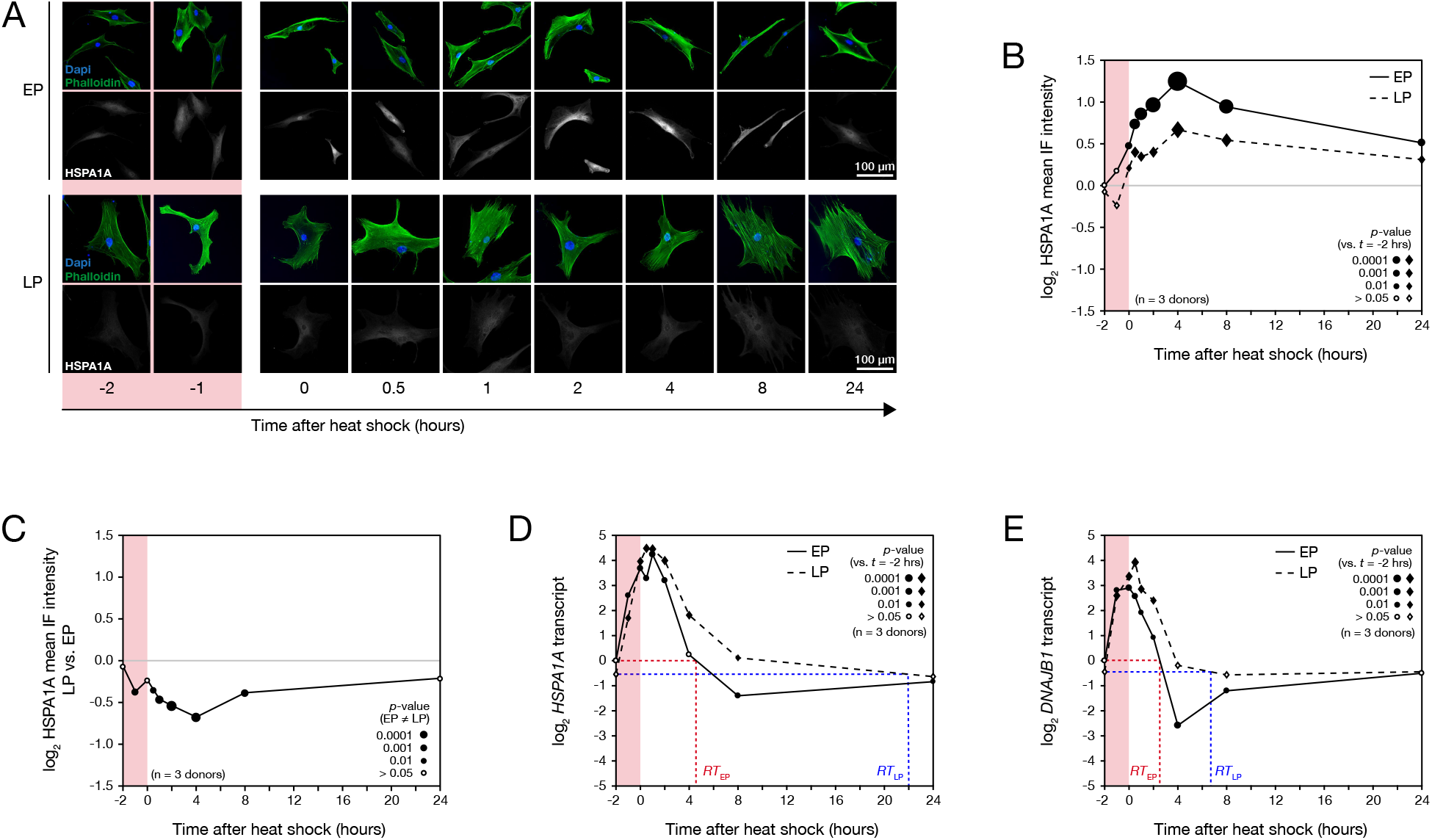
Early passage (EP) human mesenchymal stem cells (hMSCs) produce significantly more heat shock protein 70 kDa (HSPA1A) in response to heat stress than late passage (LP) cells. (**A**) Representative immunofluorescence (IF) images of heat shock protein 70 kDa (HSPA1A) in EP and LP hMSCs before, during and for 24 hours following a 2-hour heat shock treatment at 42 °C. (**B**) Quantification of mean HSPA1A intensities in images shown in panel (A). Size of points indicates significance of change in protein level vs. pre-heat shock, with solid points showing *p* < 0.05 (from ANOVA testing, *n* = 3 donors). (**C**) Ratios of mean intensities of HSPA1A in LP vs. EP hMSCs before, during and after heat shock. Point size indicates significance of where HSPA1A levels in EP ≠ LP, with solid points showing *p* < 0.05 (from ANOVA testing, *n* = 3 donors). (**D**) Corresponding levels of heat shock protein 70 kDa transcript (*HSPA1A*) in EP and LP hMSCs before, during and following heat stress. The recovery time (RT) required to return to pre-stress levels of *HSPA1A* was ∼4 hours in EP hMSCs, but 22 hours in LP cells. (**E**) Levels of DnaJ homolog subfamily B member 1 transcript (*DNAJB1*). The RT to pre-stress levels was ∼2 hours in EP hMSCs, versus ∼6 hours in LP. In panels (D) and (E), point size indicates significance of change in transcript level vs. pre-heat shock, with solid points showing *p* < 0.05 (from ANOVA testing, *n* = 3 donors).

### Senescence has minimal impact on transcriptional regulation of the heat shock response

We next examined whether the failure of LP hMSCs to match the magnitude of the HSPA1A stress response in EP cells was due to supressed levels of transcript. However, quantification of *HSPA1A* transcript by RT-qPCR showed the response in EP and LP hMSCs to be remarkably similar: starting from comparable pre-stress levels, in both cases transcripts rose rapidly during heat treatment, and reached a similar maximum level an hour into the recovery period (Fig. 4D). The primary difference between the behaviour of EP and LP cells was the time required to return to the pre-stress transcript level (recovery time *RT*_*EP*_ vs. *RT*_*LP*_, indicated on Fig. 4D): in EP cells, *HSPA1A* was insignificantly different from the initial level after 4 hours of recovery; in LP cells, this recovery time was greater than 8 hours (significance from ANOVA indicated on Fig. 4D). The time difference between peaks in *HSPA1A* transcript and levels of the encoded protein was about 3 hours (comparing Figs. 4B and D), consistent with earlier characterisations of the heat stress response in yeast (Jarnuczak et al, 2018). This delay implies that an initial regulation of HSPA1A must be post-translational. We also quantified the stress response of *DNAJB1*, encoding the HSP70 cofactor DnaJ homolog subfamily B member 1, a member of the HSP40 family (Hartl et al, 2011). The results here mirrored those of *HSPA1A*: a rapid initial rise from similar levels in EP and LP hMSCs, reaching a maximum an hour into the recovery period, before a return to baseline levels; we also again found that *RT*_*EP*_ < *RT*_*LP*_ (Fig. 4E). The stress response is understood to be regulated by the transcription factor HSF1 (Jayaraj et al, 2020). In the absence of stress, HSF1 forms a complex with chaperone proteins located in the cytoplasm. Stress conditions cause this complex to disassemble, releasing HSF1 and enabling it to translocate to the nucleus where it can activate the transcription of stress response proteins. Our data suggests that this mechanism is unaffected by senescence: the rapidity and magnitude of the transcriptional response is similar in LP vs. EP hMSCs. In support of this, quantification of HSF1 protein in the nucleus by IF showed no significant differences between EP and LP hMSCs following heat stress (Supplemental Figs. S4E-G). In summary, senescent hMSCs appeared to misregulate their stress response at a protein level, rather than through an inability to control transcription. We would therefore interpret the slower return of chaperone transcripts to baseline levels in senescent cells following stress as evidence that the burden of unfolded protein persists for longer.

### Loss of protein turnover and translation machineries are limiting factors in the stress response of senescent cells

To investigate how senescence affects the proteotoxic stress response, and predict the consequence of any alterations, we utilised a system of ordinary/delay differential equations (ODE/DDE) to model the concentrations of key factors, using parameters informed by our experimental findings. Building on previous models (Sivery et al, 2016; Zheng et al, 2016), our model considered interactions between four populations: the chaperone protein HSPA1A, the transcription factor HSF1, the E3 ubiquitin-protein ligase CHIP (an abbreviation of ‘C-terminus of HSP70 interacting protein’, and protein product of the *STUB1* gene), and intracellular misfolded protein (MFP). Reaction rates were derived to represent seven well-defined biological processes between these populations (Fig. 5A; Supplemental Table S6). CHIP was included in the model following our observation that short-term regulation of HSPA1A was principally imposed downstream of transcription. CHIP is known to work with HSP70 and HSP90 family proteins in the HSR, but is also a cofactor in HSPA1A turnover (Kundrat & Regan, 2010; Tawo et al, 2017; Zhang et al, 2015). CHIP has been shown to ubiquitinate HSPA1A and its client proteins, marking them for turnover, but with greater affinity for the misfolded protein than for the chaperone (Qian et al, 2006). This mechanism reduces the turnover rate of chaperones while the concentration of misfolded protein is high, and increases the turnover rate of chaperones during recovery from stress, where misfolded protein concentration is low, but chaperone concentration is elevated.

**Figure 5.**
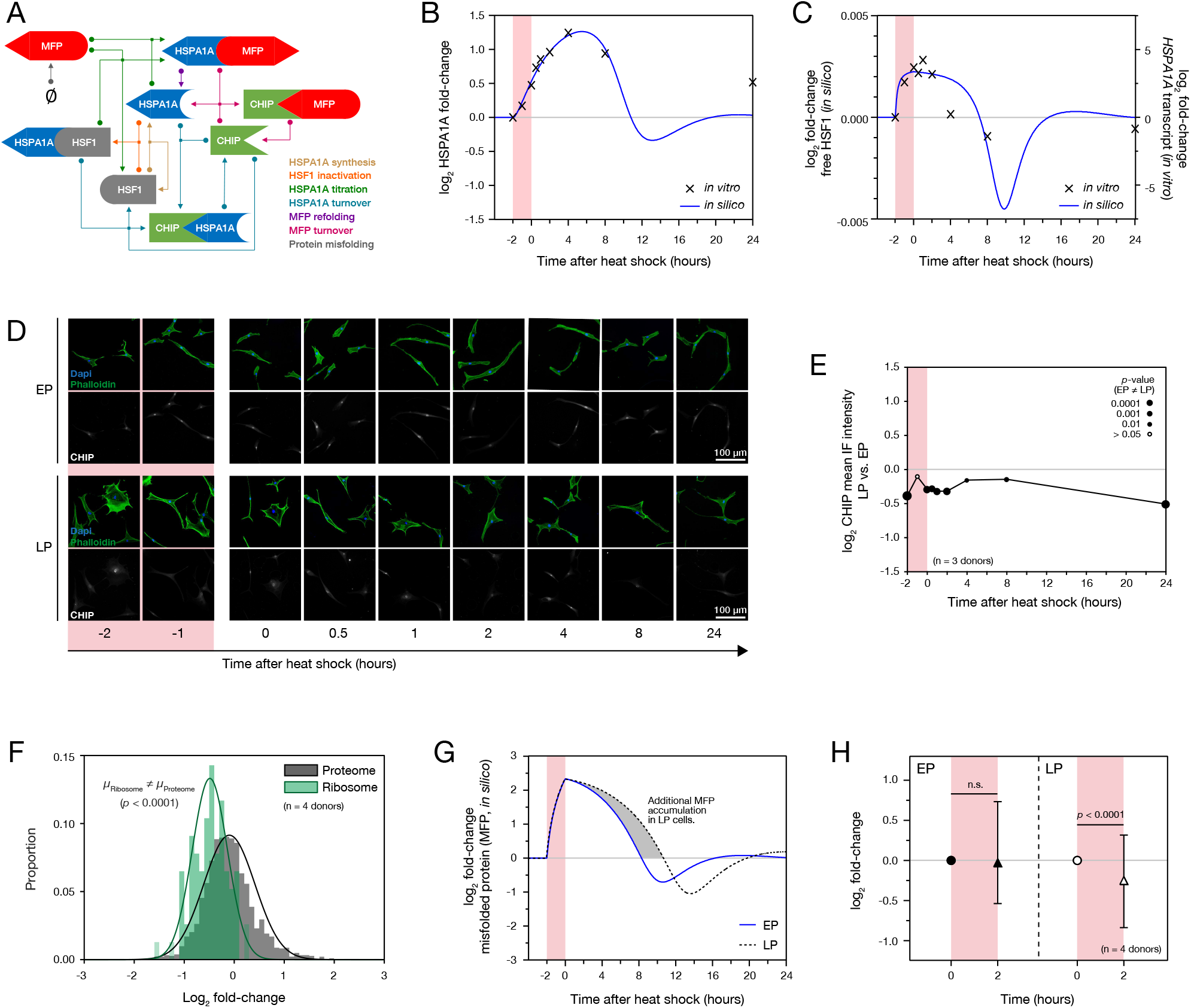
Loss of translational capacity and misfolded protein ubiquitination machinery causes increased proteotoxicity in senescent cells. (**A**) Schematic of the titration model of HSPA1A regulation. Each arrow colour represents a process with rates defined in the Materials and Methods section and Supplementary Table S6. Arrows begin at the process reactants and end at the process products. (**B**) Ordinary / delayed differential equation (ODE/DDE) model simulation of the stress response. The *in silico* model was given sufficient time to reach a stable equilibrium, before a proteotoxic stress was simulated by increasing the rate at which misfolded proteins were generated within the model for 120 minutes. At each time interval, the *in silico* concentration of HSPA1A was recorded, and is shown overlaid with data acquired by IF from EP hMSCs *in vitro* (Fig. 4B). (**C**) As (B), but showing concentrations of modelled active HSF1 overlaid, with a scaling factor, onto experimentally-derived *HSPA1A* transcript levels in EP hMSCs (Fig. 4D). (**D**) Representative immunofluorescence (IF) images of E3 ubiquitin-protein ligase CHIP in early passage (EP) and late passage (LP) primary human mesenchymal stem cells (hMSCs) before, during and for 24 hours following a 2-hour heat shock treatment at 42 °C. (**E**) Ratios of intensities of CHIP in LP vs. EP hMSCs before, during and after heat shock. Point size indicates significance of where EP ≠ LP, with solid points showing *p* < 0.05 (from ANOVA testing, *n* = 3 donors). (**F**) Levels of proteins identified as constitutive of the ribosome (77 proteins) compared to the whole proteome in LP vs. EP hMSCs in the absence of heat shock. Curves show Gaussian fits to the data (with maxima at *µ*_Ribosome_ and *µ*_Proteome_). Levels of ribosomal proteins were supressed relative to the whole proteome in LP vs. EP cells (*p* < 0.0001, two-tailed t-test). (**G**) ODE/DDE model simulation of misfolded protein levels in the stress response of EP and LP cells. (**H**) The distribution (median and interquartile range) of abundance changes of 359 monobromobimane (mBBr)-tagged peptides from 252 proteins in response to stress in EP and LP hMSCs subjected to 2-hour heat treatment at 42 °C. Significance calculated from non-parametric Wilcoxon signed-rank test, *n* = 4 donors.

The model was used to simulate a 2-hour heat shock, comparing the dynamics of HSPA1A levels calculated *in silico* to those acquired experimentally in EP cells (Fig. 5B). A test of accuracy was performed by comparing *in silico* and experimental half-lives of HSPA1A: switching off HSPA1A synthesis in the model resulted in a half-life of 100 minutes (Supplemental Fig. S5A), comparing well with a literature value of 99 minutes (Mao et al, 2013). Furthermore, if the model accurately described HSF1 activation dynamics, we would expect it to closely follow the dynamics of *HSPA1A* transcript, with an associated scaling factor. To investigate this relationship, we superimposed active HSF1 concentrations evaluated from *in vitro* modelling with experimentally derived *HSPA1A* mRNA levels in EP cells (Fig. 5C). Active HSF1 increased immediately following stress, reaching a plateau after approximately one hour; active HSF1 then gradually decreased, dropping below the levels observed in the pre-stress equilibrium state before recovering to initial levels. This behaviour was matched by our experimental measurements of *HSPA1A* mRNA.

Having verified the consistency of our ODE/DDE model with findings *in vitro*, we sought to investigate how changes observed in senescent cells could affect simulations of the stress response. With our earlier experimental observations of the temporal dynamics of the HSPA1A stress response (Figs. 4A-D) supporting the hypothesis that transcription takes hours to dominate HSPA1A regulation, CHIP concentrations in EP and LP cells were analysed to investigate whether a change in CHIP-mediated turnover could be a limiting factor in the senescent stress response. IF imaging showed a significant decrease in CHIP levels in EP vs. LP cells (Fig. 5D; see Fig. 5E for significance values, determined by ANOVA). In contrast, senescence had a minimal effect on levels of transcription factor HSF1 (Supplemental Fig. S4G). Following our finding that the limiting factor in the stress response of LP cells was downstream of chaperone transcription, we performed a relative quantification of the levels of ribosomal proteins in our EP vs. LP mass spectrometry data. A comparison of quantities of 77 detected ribosomal proteins showed a significant decrease in ribosomal machinery in senescence (a mean log_2_ fold-change of −0.4873; *p* < 0.0001, two-tailed t-test; Fig. 5F), suggestive of a loss of translational capacity.

### In silico and experimental measures of misfolded protein suggest senescent cells are more susceptible to accumulation of proteotoxic damage

Our experimental observations of senescence-induced loss of CHIP and translational capacity were incorporated into our ODE/DDE model by (i) multiplying the pre-stress CHIP concentration by a factor of 2^− 0.7799^ = 0.5824 (Fig. 5E); and, (ii) multiplying the parameter *k* – representing the maximal rate of HSPA1A synthesis – by a factor of 2^− 0.4873^ = 0.7134 (Fig. 5F). *In silico*, these alterations lead to a decrease in the magnitude of HSPA1A upregulation in response to stress, and an extension of the time taken for transcriptionally active HSF1 to return to pre-stress levels (Supplementary Figs. S5B, C) – both consistent with our *in vitro* quantification of HSPA1A protein and transcript in the stress responses of LP cells. The model enabled us to estimate the increased burden of misfolded proteins in senescent cells subjected to heat stress – a quantity difficult to measure directly by experimental means (Fig. 5F): the period of proteotoxic stress was prolonged in senescent cells (from 10.3 to 12.7 hours), and the area under the MFP curve (i.e. persistence of misfolded protein in the cell) increased by approximately 30%.

We tested for an increased presence of proteins with altered folding states experimentally using mass spectrometry in conjunction with monobromobimane (mBBr) protein labelling (Gilbert et al, 2019; Johnson et al, 2007; Swift et al, 2013). mBBr selectively labels reduced cysteine residues typically buried in the hydrophobic cores of folded proteins, however reactivity can be modulated by protein unfolding (increasing access of mBBr to cysteine residues) or aggregation (decreasing access). As such, modulated levels of mBBr-adduct peptides quantified by LC-MS/MS, in comparison to a control state, can be interpreted as a change to the folding-state of the proteome. Here, a broadening of the distribution of abundances of mBBr-adduct peptides indicated protein conformational changes in response to thermal stress (Supplemental Fig. S5D). Figure 5H shows mean log_2_ fold-changes to 359 unique mBBr-adduct peptides from 252 proteins detected in EP and LP hMSCs subjected to a 2-hour treatment at 42 °C. Consistent with our *in silico* prediction that proteostasis would be perturbed to a greater extent in LP cells under heat stress, a significant change was observed in the mBBr-labelling profiles of LP cells (*p* < 0.0001, t-test), but no significant change was found in EP cells. Together, the ODE/DDE model and mBBr labelling experiment show that a consequence of an abrogated stress response is that senescent cells are more liable to the accumulation of proteotoxic damage.

## DISCUSSION

Mathematical models of biological processes have increasingly been used not only to validate hypotheses, but also as means to generate new hypotheses (Gunawardena, 2014; Tomlin & Axelrod, 2007). Biological processes are highly multifaceted, and a functional descriptive model can demonstrate a working understanding of a process. Going further, one can perturb single elements of a system *in silico*, or make measurements or predictions about system features that would be difficult to measure directly via experiment. Here, we created an ODE/DDE model of the stress response with the aim – when informed by temporally-resolved experimental data – of challenging the current understanding of HSPA1A regulation. Models attempting to describe this process have thus far been built around the theory of mass synthesis of an already highly-abundant protein, and have generally neglected post-translational regulation (Magni et al, 2018; Pal & Sharma, 2020; Petre et al, 2011; Rieger et al, 2005; Scheff et al, 2015; Sivery et al, 2016; Szymanska & Zylicz, 2009; Zheng et al, 2016). Our model has built on the minimal models developed by Sivery et al. and Zheng et al. The former of these models assumed an unchanged rate of HSPA1A turnover during stress, while the latter disregarded HSPA1A turnover altogether. Both designs therefore neglected evidence of upregulated HSPA1A half-life during stress (Li & Duncan, 1995; Mao et al, 2013; Qian et al, 2006). Moreover, predominantly stabilising chaperone machinery under stress, rather than synthesising additional protein, is supported in the literature (Chen et al, 2015; Mao et al, 2013). This suggests an important general point: that when faced with a need to rapidly remodel protein machinery in order to address an acute or short-term demand, particularly in high number-density proteins such as those of the chaperome or cytoskeleton, cells may draw on other tools besides an upregulation of translation. It may in many cases be faster, and more efficient in terms of conserving cellular resources, to regulate post-translational modifications and turnover rates (and potentially also location within the cell, as evidenced by our imaging data). These factors may also be of key importance to understanding the consequences of cellular senescence, and by extension how cells such as MSCs might be used in therapy, as well as broader facets of the ageing process and age-associated disease. Our *in silico* simulations have predicted that senescent-associated loss of translational and ubiquitin-ligase capacities leads to a greater accumulation of misfolded proteins during proteotoxic stress. Using a cysteine-reactive label which binds to residues typically buried in correctly folded proteins, this prediction was substantiated by showing significant conformational changes to the proteomes of senescent cells in response to stress, whereas no perturbation was seen in proliferating cell populations.

## Materials and Methods

### Primary cell culture

Human mesenchymal stem cells (hMSCs) were isolated from bone marrow using established methodology (Strassburg et al, 2010). All experiments were performed in accordance with relevant guidelines and regulations, and with National Research Ethics Service and University of Manchester approval. hMSCs were cultured on tissue culture treated polystyrene (TCTP) in low-glucose DMEM with pyruvate (Thermo Fisher Scientific) supplemented with 10% fetal bovine serum (FBS, Labtech.com) and 1% penicillin/streptomycin cocktail (PS, Sigma-Aldrich). Details of individual donors and the passage numbers at which hMSCs were used are provided in Table 1.

### Heat shock and inhibitor treatments

For immunofluorescence (IF) assays, hMSCs were seeded at a density of 500 cells/cm^2^ onto 1.5 mm-thick glass coverslips (SLS) in 35 mm petri dishes (Corning) for 24 h prior to experimentation. For proteomic and transcript assays, hMSCs were seeded at a density of 7000 cells/cm^2^ into T75 flasks (Corning) 24 h prior to experimentation. Cells were incubated for 2 h at 42 °C in a 5% CO_2_ humidified incubator, while control groups were maintained at 37 °C. For IF imaging and RT-qPCR, nine time points were taken across the 2 h heat shock (HS) and 24 h recovery period: pre-HS; 1 h into HS; immediately following HS; 30 min post-HS; and 1, 2, 4, 8 and 8 h post-HS. In experiments with HSP70 inhibition, media was replaced with fresh media containing 0.01% DMSO (Sigma) and 10 µM 2-phenylethynesulfonamide (PES, Sigma) 30 min prior to HS; comparisons were made to vehicle-only controls. HSP90 inhibitors were used at concentrations of 250 nM for geldanamycin (Sigma) (Cheung et al, 2010) and 1 µM for radicicol (Sigma) (Schulte et al, 1999), delivered in fresh media containing 0.01% DMSO; cells were treated for 24 h, before comparison to vehicle-only controls (note that direct comparisons were not made between HSP70 and HSP90 inhibition datasets).

### Immunofluorescence (IF), microscopy and image analysis

Human MSCs were fixed with 4% paraformaldehyde (PFA, VWR International) in deionized (DI) water for 10 min at 37 °C, followed by washing in Dulbecco’s phosphate-buffered saline (PBS, Sigma). Cells were permeabilized in 1% Triton-X (Sigma-Aldrich) in PBS for 10 min and blocked with 2% bovine serum albumin (BSA, Sigma-Aldrich), 0.1% Triton-X in PBS at 37 °C for 1 h. Samples were incubated with a monoclonal antibody raised in rabbit against HSPA1A (1:1000; Abcam, ab181606) for 1 h at 37 °C, followed by 4 x washes in PBS. The secondary AlexaFluor-594 donkey anti-rabbit (1:1000; ThermoFisher Scientific, A21207) was added with DAPI (1:500; Sigma Aldrich, D9542) and AlexaFluor-488 Phalloidin (1:500; Cell Signaling Technology, #8878), incubated for 1 h and washed 5 x times with PBS. Coverslips were rinsed 2 x in DI water before mounting onto 1 mm-thick glass slides (Thermo Scientific) using anti-fade mounting medium (Dako).

Images were collected on a Zeiss Axioimager.D2 upright microscope using 10x and 20x / 0.5 EC Plan Neofluar objective lenses and captured using a Coolsnap HQ2 camera (Photometrics) with Micro-Manager software (version 1.4.23). Band pass filter sets for DAPI, FITC and Texas red were used to prevent bleed between channels. Images were processed using Fiji and ImageJ (version 2.0.0, National Institutes of Health, USA); CellProfiler (version 2.1.1, Broad Institute, USA) (Kamentsky et al, 2011) was used to quantify cell morphometric parameters. Images were corrected for background fluorescence by subtracting the mean intensity/pixel of a cell-free area from each pixel; all images under comparison in the same experiment had matched exposure and contrast settings.

### RT-qPCR

MSCs were harvested with EDTA-trypsin (Sigma) and pelleted by centrifugation. RNA was extracted from cell pellets using the RNeasy Mini kit (Qiagen), as per the manufacturer’s instructions, and its concentration was measured using a NanoDrop 2000 spectrophotometer (Thermo Fisher). 1 μg of mRNA per sample was reverse transcribed using the High Capacity RNA-to-cDNA Kit (ThermoFisher Scientific) in a Verity Thermal Cycler (Applied Biosystems). RT-qPCR was performed in triplicate using SYBR Select Master Mix (ThermoFisher Scientific) using a StepOnePlus Real-Time PCR System (ThermoFisher Scientific). Data was analysed using the 2^−ΔΔCt^ method (Livak & Schmittgen, 2001) and normalised to the housekeeping gene *PPIA*. Custom designed and validated primers (PrimerDesign Ltd) were used as follows:

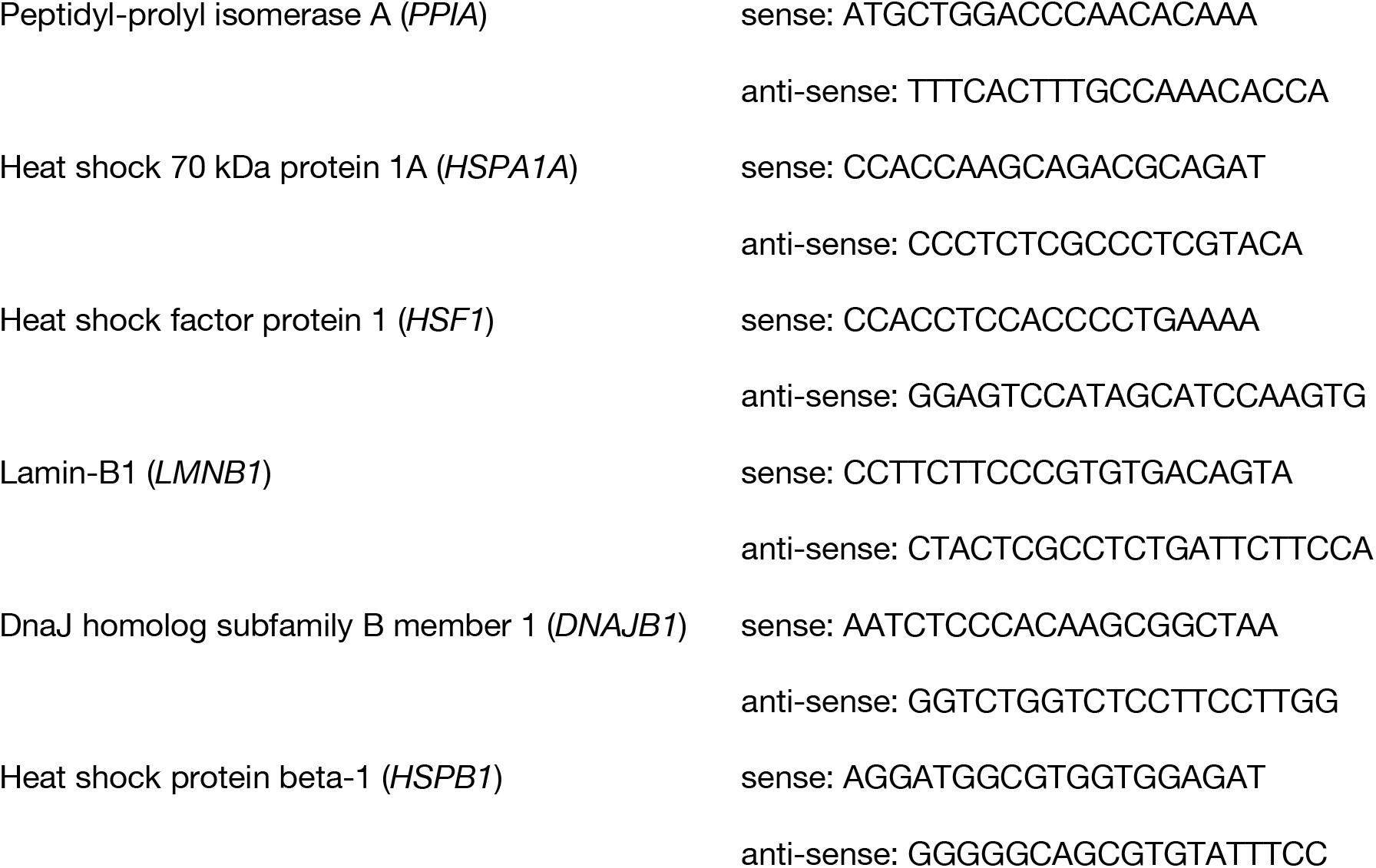

### Mass spectrometry (MS) sample preparation and analysis

hMSCs were washed with PBS, harvested with EDTA-trypsin (Sigma), pelleted by centrifugation, re-suspended in PBS and re-pelleted into 1.5 mL LoBind tubes (Eppendorf). Six 1.6 mm steel beads (Next Advance) were added to the cell pellet tube with 30 µL SL-DOC (1.1% sodium dodecyl sulfate (Sigma), 0.3% sodium deoxycholate (Sigma), 25 mM ammonium bicarbonate (AB, Fluka), 0.1% protease inhibitor cocktail set 1 (Calbiochem) and 0.1% phosphatase inhibitor no. 3 (Sigma) in de-ionised (DI) water). Cells were homogenized in a Bullet Blender (Next Advance) at maximum speed for 2 min. Homogenates were cleared by centrifugation (20k g, 5 min). Protein concentrations were measured by Direct Detect spectrophotometer (Millipore). Immobilized-trypsin beads (Perfinity Biosciences) were suspended in 200 µL of digest buffer (1 mM CaCl_2_ (Sigma) in 25 mM AB) containing 50 µg of lysate protein and shaken overnight in a thermomixer (Eppendorf; 1400 rpm at 37 °C).

The resulting digest was reduced (addition of 4 µL x 0.5 M dithiothretol (Sigma) in 25 mM AB; 10 min shaking at 60 °C) and alkylated (addition of 12 µL x 0.5 M iodoacetamide (Sigma) in 25 mM AB; 30 min shaking in the dark at RT). Peptides were acidified by addition of 5 µL x 10% trifluoroacetic acid (Riedel-de Haën) in DI water, and cleaned by two-phase extraction (2 x addition of 200 µL ethyl acetate (Sigma) followed by vortexing, centrifugation and aspiration of the organic layer). Peptides were desalted using POROS R3 beads (Thermo Fisher), in accordance with the manufacturer’s protocol, and lyophilized. Peptide concentrations (measured by Direct Detect spectrophotometer, Millipore) in injection buffer (5% HPLC grade acetonitrile (ACN, Fisher Scientific) 0.1% trifluoroacetic acid in DI water) were adjusted to 0.3 g/L.

Samples were analysed by LC-MS/MS using an UltiMate® 3000 RSLC (Dionex Corporation) coupled to a Q Exactive HF (Thermo Fisher Scientific) mass spectrometer. Peptides were separated using a 75 mm × 250 μm inner diameter 1.7 μM CSH C18 analytical column (Waters) with a gradient from 95% A (0.1% formic acid, FA, Sigma) and 5% B (0.1% FA in ACN) to 7% B at 1 min, 18% B at 58 min, 27% B in 72 min, and 60% B at 74 minutes at 300 nL/min. Peptides were selected for fragmentation automatically by data-dependent analysis.

### Proteomics data processing

Alignment and peak-picking were performed in Progenesis QI (Waters) and searched using Mascot (Matrix Science UK), against the SWISS-Prot and TREMBL human databases. The peptide database was modified to search for alkylated cysteine residues (monoisotopic mass change, 57.021 Da), oxidized methionine (15.995 Da), hydroxylation of asparagine, aspartic acid, proline or lysine (15.995 Da) and phosphorylation of serine, tyrosine, threonine, histidine or aspartate (79.966 Da). A maximum of 2 missed cleavages was allowed; peptides with charges above +4 or fewer than two isotopes were removed. Peptide detection intensities were exported from Progenesis QI as Excel spreadsheets (Microsoft) for further processing.

Proteomics datasets were analysed using code written in-house in MATLAB with the bioinformatics toolbox (R2015a, The MathWorks, USA). Raw ion intensities from peptides from proteins with fewer than 3 unique peptides per protein were excluded from quantification. Peptide lists were filtered leaving only those peptides with a Mascot score corresponding to a Benjamini-Hochberg false discovery rate (BH-FDR) (Benjamini & Hochberg, 1995) of < 0.2. Normalisation was performed as follows: raw peptide ion intensities were log-transformed to ensure a normal distribution and normalised within-sample by equalising sample medians (subtracting sample median). Fold-change differences in the quantity of proteins detected in different samples were calculated by fitting a linear regression model that takes into account donor variability at both the peptide and protein levels (Gilbert et al, 2019; Mallikarjun et al, 2020).

### Reactome graphs

A pathway representation analysis of protein-level fold-changes (Fig. 1E) was carried out using the Reactome Pathway Analysis tool (Pathway browser version 3.6, Reactome database release 71) (Fabregat et al, 2017; Fabregat et al, 2018; Jassal et al, 2020). Significantly represented pathways at the FDR-corrected 5% level were coloured according to the mean log_2_-fold change of proteins within the pathway.

### Modularity analysis

The 332 protein human chaperome (Brehme et al, 2014) was modelled as a weighted undirected network in order to discern its community structure (Fig. 2A, Supplementary Fig. S2A). Chaperones with at least one interaction satisfying the highest confidence filter from STRING database (version 11) (Szklarczyk et al, 2019) were used as the nodes of the network, whilst these highest confidence interactions between chaperones were used as undirected edges in the network. Edges between nodes were weighted according to the STRING interaction score between the respective chaperones. The network modularity, *Q*^*W*^*ϵ* [−0.5,1], was used to give a measure of how well the network separated into non-overlapping communities of highly-interconnected nodes. A network with edges distributed at random would score a modularity of zero, indicating no presence of community structure, whilst higher scores would indicate more intramodular edges than would be expected at random. The maximal modularity score was calculated using methods described previously (Newman, 2006; Rubinov & Sporns, 2010):

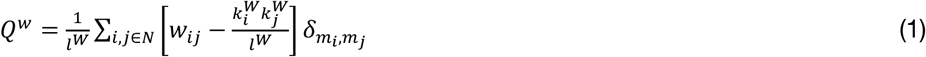

Where *i* and *j* denote two elements of the set of all nodes *N*, connected by an edge of weight *w*_*ij*_*ϵ*(0,1]. *l*^*W*^ = ∑_*i,j∈N*_ *w*_*ij*_ is the sum of all weights in the network; 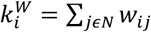 is the weighted degree of node *i*; and 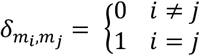 is the Kronecker delta function, where *m*_*i*_ is the module containing node *i*. 942 interactions were identified between 181 chaperone proteins. Supplementary Fig. S2B shows the adjacency matrix *A*, whose elements *A*_*ij*_ are the weights of edges between two elements *i, j* ∈ *N*. To calculate the modularity of the network, *A* was used as the input for the Matlab Brain Connectivity Toolbox (Rubinov & Sporns, 2010). The maximal modularity of the human chaperome was found to be *Q*^*w*^ = 0.5328, with a community structure consisting of 19 modules. The 5 modules with the most proteins seen in our label-free mass spectrometry dataset were investigated and named according to the function of chaperones within the module.

### Linear modelling

Linear regression models were fit to data in MATLAB using the *fitlm* function (R2015a, MathWorks) using the equation:

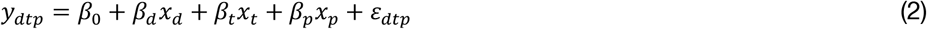

The response variable *y*_*dtp*_ (e.g. HSPA1A concentration) was modelled as being dependent on donor *d*, temperature *t* (categorical variable, either with or without heat shock), and passage number *p* (categorical variable, either early passage (EP) or late passage (LP, i.e. senescent), plus an error term, *ε*_*dtp*_.

### Rate equations used to describe the heat-stress response

We established a model of the heat-stress response that considered the concentrations of six key species: HSF1 (active HSF1 protein), HSPA1A, CHIP (E3 ubiquitin-protein ligase CHIP protein), MFP (misfolded protein), HSPA1A-HSF1 (HSPA1A in complex with inactive HSF1), and HSPA1A-MFP (HSPA1A bound to a misfolded protein). The model was based on the following seven reactions, with reaction rates *k*_*i*_ (Fig. 5A):

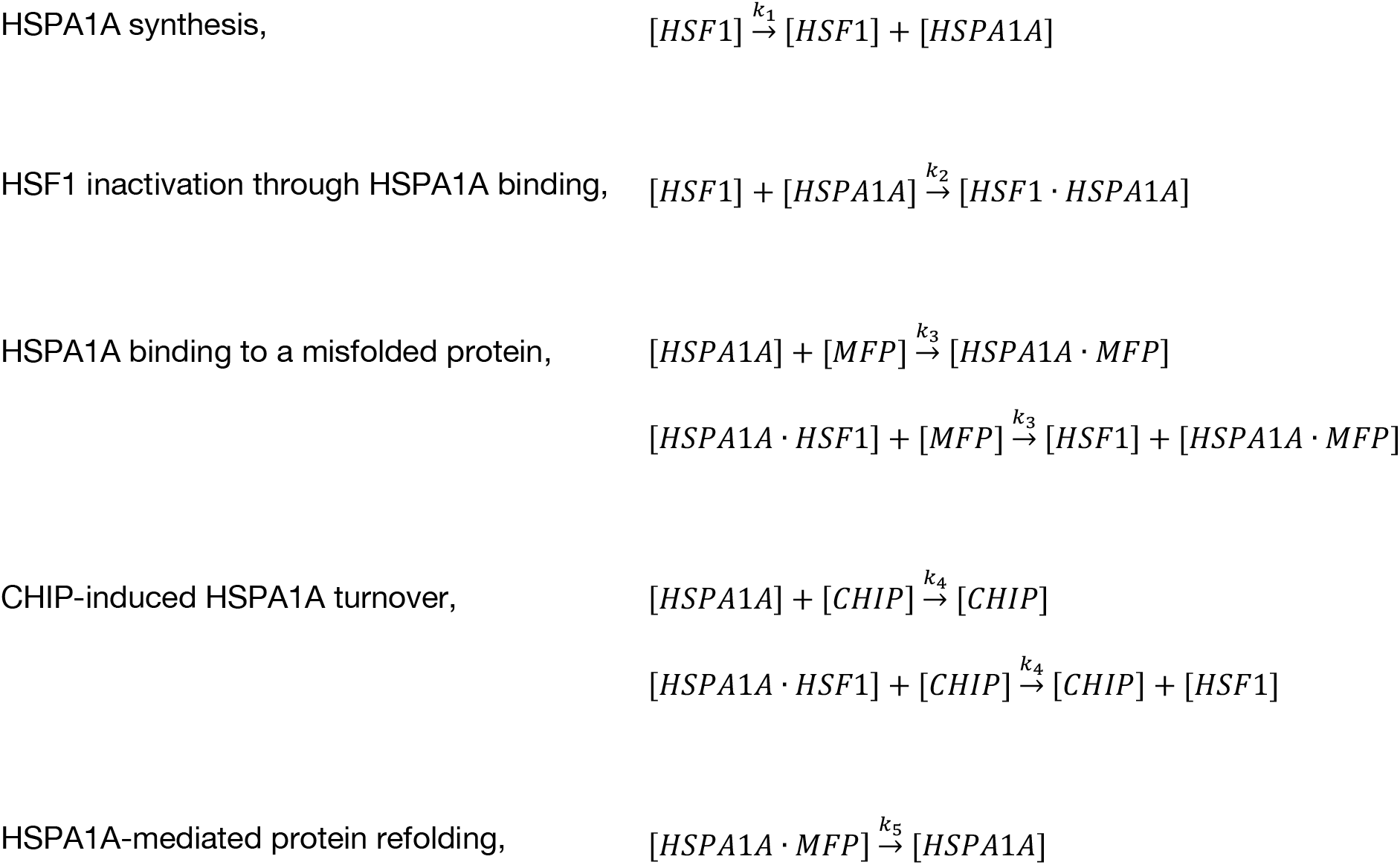
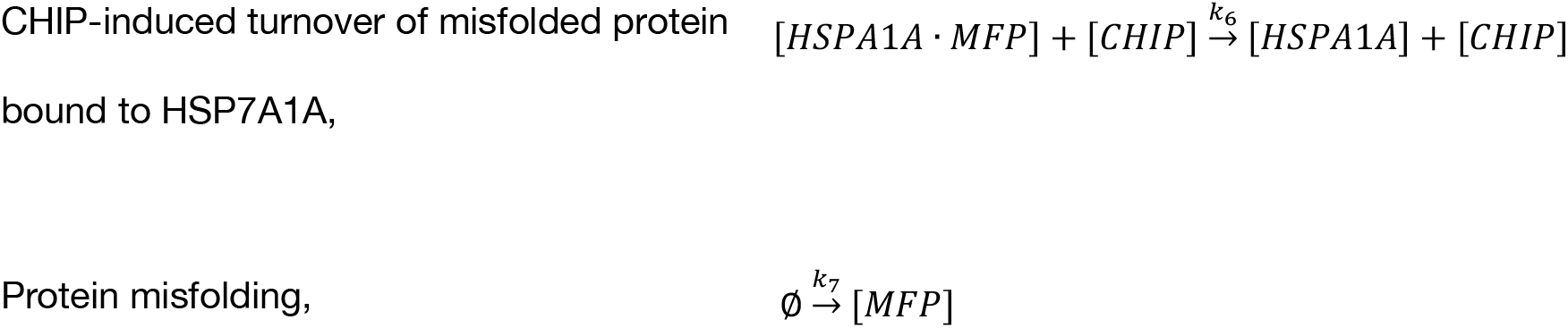

HSPA1A synthesis was modelled using a Hill function,

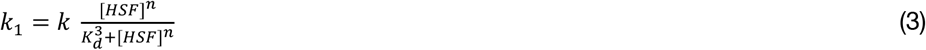

Where the Hill coefficient, *n* = 3, was chosen to reflect HSF1 trimerisation; 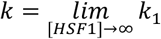 was the maximal transcription rate and *K*_*d*_ the HSF1 dissociation constant.

### Initial conditions, constraints and running of the model

Initial conditions and constraints were based on experimental and literature-sourced results. Concentrations of component species were set relative to HSF1, quasi-arbitrarily set at 0.1 µM, based on previous estimates (Milo, 2013). The mean mass spectrometry signal ratio between HSPA1A and the E3 ubiquitin-protein ligase CHIP across the four early passage (EP), pre-stress biological replicates (Supplementary Fig. S5F), was used as the ratio of concentrations between HSPA1A and CHIP at equilibrium. Practically, this was done by adjusting CHIP levels based on HSPA1A at every time point until a stable equilibrium was reached (Supplementary Fig. S5E). It was assumed that there would be no substantial difference between the binding affinity of HSPA1A to HSF1 or client proteins, such that *k*_2_ ≈ *k*_3_. The turnover time for a protein via the proteasome was assumed to be linearly proportional to its length in amino acids. To incorporate this, the ratio between the number of amino acids in HSPA1A and the median length of a human protein (Brocchieri & Karlin, 2005) was used to estimate the relative time taken for turnover following ubiquitination, *k*_6_: *k*_4_ = 641: 375. Parameters that could not be inferred from theory, directly from experimental results, or literature evidence (*k, K*_*d*_, *k*_2_, *k*_4_, *k*_5,_ *k*_7_) were chosen to optimise the model’s ability to replicate *in vitro* HSPA1A protein dynamics observed in IF data (Figs. 4A, B). This was done by using the *MATLAB fminsearch* function to minimise the total sum of squared errors (SSE) between *in vitro* measurements of HSPA1A concentration *and in silico* values calculated at matched timepoints. Several orders of magnitude were trialled for each remaining variable to obtain initial conditions for the optimisation (Supplementary Figs. S5H-I). A summary of all parameters can be found in Supplemental Table S6.

The temporal dynamics of the heat shock response were modelled using MATLAB (R2015a, MathWorks). All populations were recalculated at discrete time intervals using ordinary differential equations (ODE), except for *HSP*70 which was calculated using a delay differential equation (DDE). This accounted for the time delay (*τ* ∼ *U*[60, 180] minutes) between transcription and translation (Jarnuczak et al, 2018). The DDE/ODEs were evaluated every (simulated) 0.01 minutes. A proteotoxic stress comparable with heat shock was simulated by multiplying the reaction rate *k*_7_ by another optimised parameter, *α* > 1 for 120 minutes.

### Protein labeling with monobromobimane (mBBr)

Media was removed from cells in T75 flasks (Corning) either pre-stress or immediately post-stress and cells were washed with PBS. Cells were then labelled by incubation with 5 mL of 400 µM monobromobimane (mBBr; Sigma-Aldrich) in PBS at 37 °C for 10 min. Following labelling, 5 mL of 2 mM glutathione in PBS was added to quench the reaction. The quenched mBBr solution was removed and cells washed with PBS, before harvesting with trypsin for MS analysis, as described above. When searching MS data, the peptide database was modified to search for mBBr adducts to cysteine (monoisotopic mass changes, 133.053 and 150.056 Da). mBBr-labelled peptides were filtered to only include sequences from reviewed protein annotations (Bateman et al, 2019) and labelled peptides with missed cleavages were summed with their fully tryptic counterparts. 359 unique mBBr-labelled peptides were detected. The log_2_ fold-change in labelling across samples was normalised to the respective log_2_ fold-change in protein abundance. A Wilcoxon signed-rank test was used to determine whether the mBBr-labelling profile changed between samples at the 95% confidence level.

## ACKNOWLEDGEMENTS

JL was supported by a Wellcome Trust Quantitative and Biophysical Biology studentship. HTJG and JS were funded by a Biotechnology and Biological Sciences Research Council (BBSRC) David Phillips Fellowship (BB/L024551/1). VM was partially supported by a studentship from the Sir Richard Stapley Educational Trust. Proteomics was carried out at the Wellcome Centre for Cell-Matrix Research (WCCMR; 203128/Z/16/Z) Biological Mass Spectrometry Core Facility. Transcriptomics was carried out at the Genomic Technologies Core Facility. Imaging was carried out at the Bioimaging Facility (supported by BBSRC, Wellcome and the University of Manchester Strategic Fund). We thank Drs. Ronan O’Cualain, Stacey Warwood, David Knight (MS), Craig Lawless, Rachel Scholey, Julian Selley, Leo Zeef (bioinformatics), Steven Marsden, Roger Meadows and Peter March (bioimaging) for Core Facility support.

## AUTHOR CONTRIBUTIONS

Investigation, JL, VM, EA, MO and HTJG; Formal Analysis, JL, VM, EA, MO, HTJG, SJH and JS; Writing – Original Draft, JL; Writing – Visualization, Review & Editing, JL, VM, HTJG, SJH and JS; Project Administration and Funding Acquisition, SJH and JS.

## DATA AVAILABILITY

Proteomics data have been deposited to the ProteomeXchange Consortium via the PRIDE partner repository with the identifiers: (1) PXD025280, EP and LP hMSCs subjected to heat shock; (2) PXD025305, EP and LP hMSCs subjected to heat shock in combination with HSPA1A inhibition; (3) PXD026706, EP hMSCs subjected to HSP90 inhibition; (4) PXD025329, monobromobimane labelling of EP and LP hMSCs subjected to heat stress.

## REFERENCES

Ambra R, Mocchegiani E, Giacconi R, Canali R, Rinna A, Malavolta M, Virgili F (2004) Characterization of the hsp70 response in lymphoblasts from aged and centenarian subjects and differential effects of in vitro zinc supplementation. Experimental Gerontology 39: 1475–1484

Balchin D, Hayer-Hartl M, Hartl FU (2016) In vivo aspects of protein folding and quality control. Science 353: 7

Bateman A, Martin MJ, Orchard S, Magrane M, Alpi E, Bely B, Bingley M, Britto R, Bursteinas B, Busiello G, Bye-A-Jee H, Da Silva A, De Giorgi M, Dogan T, Castro LG, Garmiri P, Georghiou G, Gonzales D, Gonzales L, Hatton-Ellis E et al (2019) UniProt: a worldwide hub of protein knowledge. Nucleic Acids Research 47: D506–D515

Ben-Zvi A, Miller EA, Morimoto RI (2009) Collapse of proteostasis represents an early molecular event in Caenorhabditis elegans aging. Proceedings of the National Academy of Sciences of the United States of America 106: 14914–14919

Benjamini Y, Hochberg Y (1995) Controlling the false discovery rate -a practical and powerful approach to multiple testing. Journal of the Royal Statistical Society Series B-Methodological 57: 289–300

Brehme M, Voisine C, Rolland T, Wachi S, Soper JH, Zhu YT, Orton K, Villella A, Garza D, Vidal M, Ge H, Morimoto RI (2014) A chaperome subnetwork safeguards proteostasis in aging and neurodegenerative disease. Cell Reports 9: 1135–1150

Brocchieri L, Karlin S (2005) Protein length in eukaryotic and prokaryotic proteomes. Nucleic Acids Research 33: 3390–3400

Calderwood SK, Murshid A, Prince T (2009) The shock of aging: Molecular chaperones and the heat shock response in longevity and aging - a mini-review. Gerontology 55: 550–558

Chen HR, Song RD, Wang GH, Ding ZH, Yang CY, Zhang JW, Zeng ZH, Rubio V, Wang LC, Zu N, Weiskoff AM, Minze LJ, Jeyabal PVS, Mansour OC, Bai L, Merrick WC, Zheng S, Shi ZZ (2015) OLA1 regulates protein synthesis and integrated stress response by inhibiting eIF2 ternary complex formation. Scientific Reports 5: 17

Cheung CHA, Chen H, Cheng L, Lyu KW, Kanwar JR, Chang J (2010) Targeting Hsp90 with small molecule inhibitors induces the over-expression of the anti-apoptotic molecule, survivin, in human A549, HONE-1 and HT-29 cancer cells. Molecular Cancer 9: 77

de Magalhaes JP, Passos JF (2018) Stress, cell senescence and organismal ageing. Mechanisms of Ageing and Development 170: 2–9

de Toda IM, Vida C, Ortega E, De La Fuente M (2016) Hsp70 basal levels, a tissue marker of the rate of aging and longevity in mice. Experimental Gerontology 84: 21–28

Deschenes-Simard X, Lessard F, Gaumont-Leclerc MF, Bardeesy N, Ferbeyre G (2014) Cellular senescence and protein degradation: Breaking down cancer. Cell Cycle 13: 1840–1858

Fabregat A, Jupe S, Matthews L, Sidiropoulos K, Gillespie M, Garapati P, Haw R, Jassal B, Korninger F, May B, Milacic M, Roca CD, Rothfels K, Sevilla C, Shamovsky V, Shorser S, Varusai T, Viteri G, Weiser J, Wu G et al (2017) The Reactome pathway knowledgebase. Nucleic Acids Research: gkx1132-gkx1132

Fabregat A, Jupe S, Matthews L, Sidiropoulos K, Gillespie M, Garapati P, Haw R, Jassal B, Korninger F, May B, Milacic M, Roca CD, Rothfels K, Sevilla C, Shamovsky V, Shorser S, Varusai T, Viteri G, Weiser J, Wu GM et al (2018) The Reactome pathway knowledgebase. Nucleic Acids Research 46: D649–D655

Gilbert HTJ, Mallikarjun V, Dobre O, Jackson MR, Pedley R, Gilmore AP, Richardson SM, Swift J (2019) Nuclear decoupling is part of a rapid protein-level cellular response to high-intensity mechanical loading. Nature Communications 10: 4149–4115

Gunawardena J (2014) Models in biology: ’accurate descriptions of our pathetic thinking’. BMC Biology 12: 11

Hartl FU, Bracher A, Hayer-Hartl M (2011) Molecular chaperones in protein folding and proteostasis. Nature 475: 324–332

Hernandez-Segura A, Nehme J, Demaria M (2018) Hallmarks of cellular senescence. Trends in Cell Biology 28: 436–453

Hipp MS, Kasturi P, Hartl FU (2019) The proteostasis network and its decline in ageing. Nature Reviews Molecular Cell Biology 20: 421–435

Hipp MS, Park SH, Hartl FU (2014) Proteostasis impairment in protein-misfolding and -aggregation diseases. Trends in Cell Biology 24: 506–514

Hsu AL, Murphy CT, Kenyon C (2003) Regulation of aging and age-related disease by DAF-16 and heat-shock factor. Science 300: 1142–1145

Huang LK, Chao SP, Hu CJ (2020) Clinical trials of new drugs for Alzheimer disease. Journal of Biomedical Science 27: 13

Jarnuczak AF, Albornoz MG, Eyers CE, Grant CM, Hubbard SJ (2018) A quantitative and temporal map of proteostasis during heat shock in Saccharomyces cerevisiae. Molecular Omics 14: 37–52

Jassal B, Matthews L, Viteri G, Gong CQ, Lorente P, Fabregat A, Sidiropoulos K, Cook J, Gillespie M, Haw R, Loney F, May B, Milacic M, Rothfels K, Sevilla C, Shamovsky V, Shorser S, Varusai T, Weiser J, Wu GM et al (2020) The Reactome pathway knowledgebase. Nucleic Acids Research 48: D498–D503

Jayaraj GG, Hipp MS, Hartl FU (2020) Functional modules of the proteostasis network. Cold Spring Harbor Perspectives in Biology 12: 18

Johnson C, Tang H, Carag C, Speicher D, Discher D (2007) Forced unfolding of proteins within cells. Science 317: 663–666

Kamentsky L, Jones TR, Fraser A, Bray MA, Logan DJ, Madden KL, Ljosa V, Rueden C, Eliceiri KW, Carpenter AE (2011) Improved structure, function and compatibility for CellProfiler: modular high-throughput image analysis software. Bioinformatics 27: 1179–1180

Kampinga HH, Craig EA (2010) The HSP70 chaperone machinery: J proteins as drivers of functional specificity. Nature Reviews Molecular Cell Biology 11: 579–592

Kaushik S, Cuervo AM (2015) Proteostasis and aging. Nature Medicine 21: 1406–1415

Klaips CL, Jayaraj GG, Hartl FU (2018) Pathways of cellular proteostasis in aging and disease. Journal of Cell Biology 217: 51–63

Koga H, Kaushik S, Cuervo AM (2011) Protein homeostasis and aging: The importance of exquisite quality control. Ageing Research Reviews 10: 205–215

Kose S, Furuta M, Imamoto N (2012) Hikeshi, a nuclear import carrier for Hsp70s, protects cells from heat shock-induced nuclear damage. Cell 149: 578–589

Kundrat L, Regan L (2010) Balance between folding and degradation for Hsp90-dependent client proteins: A key role for CHIP. Biochemistry 49: 7428–7438

Labbadia J, Morimoto RI (2015) The biology of proteostasis in aging and disease. Annual Review of Biochemistry 84: 435–464

Leu JIJ, Barnoud T, Zhang G, Tian T, Wei Z, Herlyn M, Murphy ME, George DL (2017) Inhibition of stress-inducible HSP70 impairs mitochondrial proteostasis and function. Oncotarget 8: 45656–45669

Leu JIJ, Pimkina J, Frank A, Murphy ME, George DL (2009) A small molecule inhibitor of inducible heat shock protein 70. Molecular Cell 36: 15–27

Leu JIJ, Pimkina J, Pandey P, Murphy ME, George DL (2011) HSP70 inhibition by the small-molecule 2-phenylethynesulfonamide impairs protein clearance pathways in tumor cells. Molecular Cancer Research 9: 936–947

Li DX, Duncan RF (1995) Transient acquired thermotolerance in drosophila, correlated with rapid degradation of hsp70 during recovery. European Journal of Biochemistry 231: 454–465

Livak KJ, Schmittgen TD (2001) Analysis of relative gene expression data using real-time quantitative PCR and the 2(T)(-Delta Delta C) method. Methods 25: 402–408

Lopez-Otin C, Blasco MA, Partridge L, Serrano M, Kroemer G (2013) The hallmarks of aging. Cell 153: 1194–1217

Magni S, Succurro A, Skupin A, Ebenhoh O (2018) Data-driven dynamical model indicates that the heat shock response in Chlamydomonas reinhardtii is tailored to handle natural temperature variation. Journal of the Royal Society Interface 15: 11

Mallikarjun V, Richardson SM, Swift J (2020) BayesENproteomics: Bayesian elastic nets for quantification of peptidoforms in complex samples. Journal of Proteome Research 19: 2167–2184

Mao RF, Rubio V, Chen H, Bai L, Mansour OC, Shi ZZ (2013) OLA1 protects cells in heat shock by stabilizing HSP70. Cell Death & Disease 4: 12

Martinez Guimera A, Welsh C, Dalle Pezze P, Fullard N, Nelson G, Roger Mathilde F, Przyborski Stefan A, Shanley Daryl P (2017) Systems modelling ageing: from single senescent cells to simple multi-cellular models. Essays in Biochemistry 61: 369–377

McArdle A, Dillmann WH, Mestril R, Faulkner JA, Jackson MJ (2003) Overexpression of HSP70 in mouse skeletal muscle protects against muscle damage and age-related muscle dysfunction. Faseb Journal 17: 355-+

Milo R (2013) What is the total number of protein molecules per cell volume? A call to rethink some published values. Bioessays 35: 1050–1055

Morimoto RI (2008) Proteotoxic stress and inducible chaperone networks in neurodegenerative disease and aging. Genes & Development 22: 1427–1438

Newman MEJ (2006) Modularity and community structure in networks. Proceedings of the National Academy of Sciences of the United States of America 103: 8577–8582

Pal S, Sharma R (2020) A mathematical model of heat shock response to study the competition between protein folding and aggregation. BioRxiv: doi.org/10.1101/2020.1104.1113.039123

Petre I, Mizera A, Hyder CL, Meinander A, Mikhailov A, Morimoto RI, Sistonen L, Eriksson JE, Back RJ (2011) A simple mass-action model for the eukaryotic heat shock response and its mathematical validation. Natural Computing 10: 595–612

Phillip JM, Aifuwa I, Walston J, Wirtz D (2015) The mechanobiology of aging. Annual Review of Biomedical Engineering 17: 113–141

Pittenger MF, Discher DE, Peault BM, Phinney DG, Hare JM, Caplan AI (2019) Mesenchymal stem cell perspective: cell biology to clinical progress. NPJ Regenerative Medicine 4: 15

Qian SB, McDonough H, Boellmann F, Cyr DM, Patterson C (2006) CHIP-mediated stress recovery by sequential ubiquitination of substrates and Hsp70. Nature 440: 551–555

Richardson SM, Kalamegam G, Pushparaj PN, Matta C, Memic A, Khademhosseini A, Mobasheri R, Poletti FL, Hoyland JA, Mobasheri A (2016) Mesenchymal stem cells in regenerative medicine: Focus on articular cartilage and intervertebral disc regeneration. Methods 99: 69–80

Rieger TR, Morimoto RI, Hatzimanikatis V (2005) Mathematical modeling of the eukaryotic heat-shock response: Dynamics of the hsp70 promoter. Biophysical Journal 88: 1646–1658

Roe SM, Prodromou C, O’Brien R, Ladbury JE, Piper PW, Pearl LH (1999) Structural basis for inhibition of the Hsp90 molecular chaperone by the antitumor antibiotics radicicol and geldanamycin. Journal of Medicinal Chemistry 42: 260–266

Rubinov M, Sporns O (2010) Complex network measures of brain connectivity: Uses and interpretations. Neuroimage 52: 1059–1069

Saez I, Vilchez D (2014) The mechanistic links between proteasome activity, aging and age related diseases. Current Genomics 15: 38–51

Sala AJ, Bott LC, Morimoto RI (2017) Shaping proteostasis at the cellular, tissue, and organismal level. Journal of Cell Biology 216: 1231–1241

Scheff JD, Stallings JD, Reifman J, Rakesh V (2015) Mathematical modeling of the heat-shock response in HeLa cells. Biophysical Journal 109: 182–193

Schulte TW, Akinaga S, Murakata T, Agatsuma T, Sugimoto S, Nakano H, Lee YS, Simen BB, Argon Y, Felts S, Toft DO, Neckers LM, Sharma SV (1999) Interaction of radicicol with members of the heat shock protein 90 family of molecular chaperones. Molecular Endocrinology 13: 1435–1448

Shimi T, Butin-Israeli V, Adam SA, Hamanaka RB, Goldman AE, Lucas CA, Shumaker DK, Kosak ST, Chandel NS, Goldman RD (2011) The role of nuclear lamin B1 in cell proliferation and senescence. Genes & Development 25: 2579–2593

Sivery A, Courtade E, Thommen Q (2016) A minimal titration model of the mammalian dynamical heat shock response. Physical Biology 13: 13

Strassburg S, Richardson SM, Freemont AJ, Hoyland JA (2010) Co-culture induces mesenchymal stem cell differentiation and modulation of the degenerate human nucleus pulposus cell phenotype. Regenerative Medicine 5: 701–711

Swift J, Ivanovska IL, Buxboim A, Harada T, Dingal PCDP, Pinter J, Pajerowski JD, Spinler KR, Shin J-W, Tewari M, Rehfeldt F, Speicher DW, Discher DE (2013) Nuclear lamin-A scales with tissue stiffness and enhances matrix-directed differentiation. Science 341: 1240104

Szklarczyk D, Gable AL, Lyon D, Junge A, Wyder S, Huerta-Cepas J, Simonovic M, Doncheva NT, Morris JH, Bork P, Jensen LJ, Mering C (2019) STRING v11: protein-protein association networks with increased coverage, supporting functional discovery in genome-wide experimental datasets. Nucleic Acids Research 47: D607–D613

Szymanska Z, Zylicz M (2009) Mathematical modeling of heat shock protein synthesis in response to temperature change. Journal of Theoretical Biology 259: 562–569

Tawo R, Pokrzywa W, Kevei E, Akyuz ME, Balaji V, Adrian S, Hohfeld J, Hoppe T (2017) The ubiquitin ligase CHIP integrates proteostasis and aging by regulation of insulin receptor turnover. Cell 169: 470-+

Tomlin CJ, Axelrod JD (2007) Biology by numbers: mathematical modelling in developmental biology. Nature Reviews Genetics 8: 331–340

Velazquez JM, Lindquist S (1984) Hsp70 - nuclear concentration during environmental-stress and cytoplasmic storage during recovery. Cell 36: 655–662

Welch WJ, Feramisco JR (1984) Nuclear and nucleolar localization of the 72,000-dalton heat-shock protein in heat-shocked mammalian-cells. Journal of Biological Chemistry 259: 4501–4513

Zhang HQ, Amick J, Chakravarti R, Santarriaga S, Schlanger S, McGlone C, Dare M, Nix JC, Scaglione KM, Stuehr DJ, Misra S, Page RC (2015) A bipartite interaction between Hsp70 and CHIP regulates ubiquitination of chaperoned client proteins. Structure 23: 472–482

Zheng X, Krakowiak J, Patel N, Beyzavi A, Ezike J, Khalil AS, Pincus D (2016) Dynamic control of Hsf1 during heat shock by a chaperone switch and phosphorylation. eLife 5: 26

