## Supplemental Information for "Senescence inhibits the chaperone response to thermal stress"

**CONTENTS**

Supplemental figures S1 – S5 … … … … … … … … 3

Supplemental table S6 … … … … … … … … 10

Supplemental references … … … … … … … … 11

### SUPPLEMENTAL FIGURES

#### Figure S1.

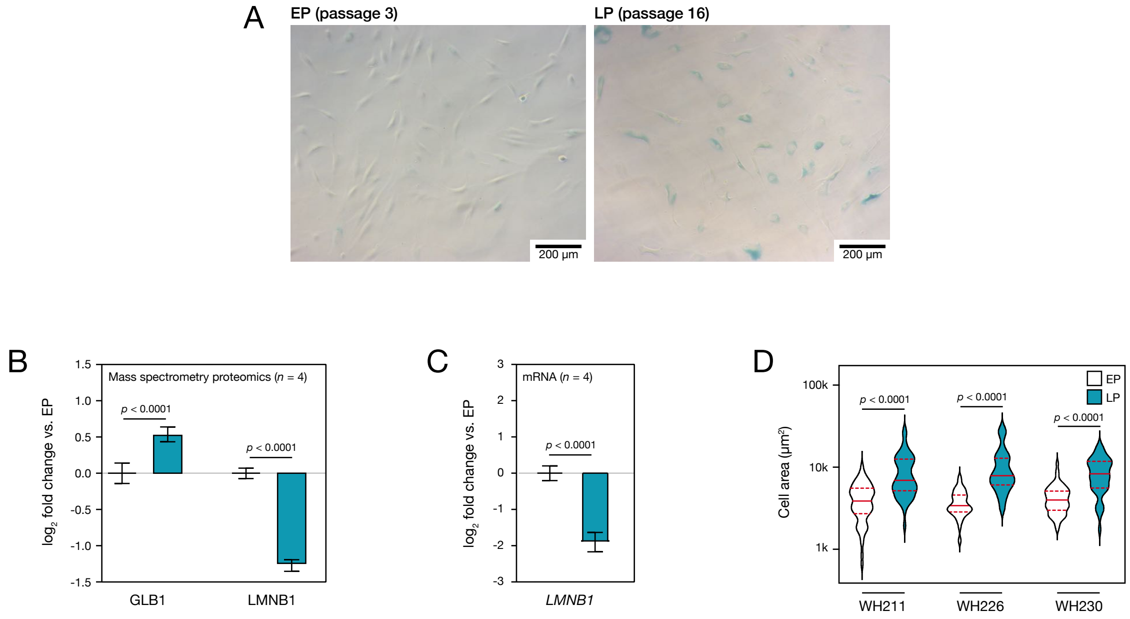


**Figure S1. Characteristics of senescence in primary human mesenchymal stem cells (hMSCs) cultured to late passage (LP) were increased relative to those in donor-matched cells at early passage (EP).** LP cells were examined at the passage number where onset of senescence occurred i.e. further cell proliferation was halted. This point was found to vary between donors (see Table 1 in the Materials and Methods section). Comparisons were made between donor-matched samples of senescent LP and EP cells at passage 3: (**A**) LP hMSCs showed positive staining for β-galactosidase. (**B**) Quantitative mass spectrometry proteomics (Fig. 1A) showed a corresponding significant increase in β-galactosidase protein (GLB1, $p < 0.0001$) in LP vs. EP cells, and significant loss of the senescence marker lamin-B1 (LMNB1, $p < 0.0001$) (Shimi et al, 2011). $\mathrm{Log}_2$ fold-changes and significance values corrected for Benjamini-Hochberg false discovery rate (BH-FDR) were calculated as described previously (Mallikarjun et al, 2020); $n = 4$ primary donors. (**C**) RT-qPCR also showed a corresponding significant decrease of *LMNB1* transcript in LP vs. EP hMSCs ($p < 0.0001$ established by t-test; normalisation to housekeeping gene *PPIA*; $n = 4$ primary donors). (D) An analysis of cell morphology showed that LP hMSCs had significantly larger spread areas than EP cells ($p < 0.0001$ established by ANOVA testing, with ≥ 39 cells imaged per condition), consistent with established senescence phenotype (Hernandez-Segura et al, 2018). Solid red lines indicate medians; dashed red lines indicate quartiles.

**Figure S2.**

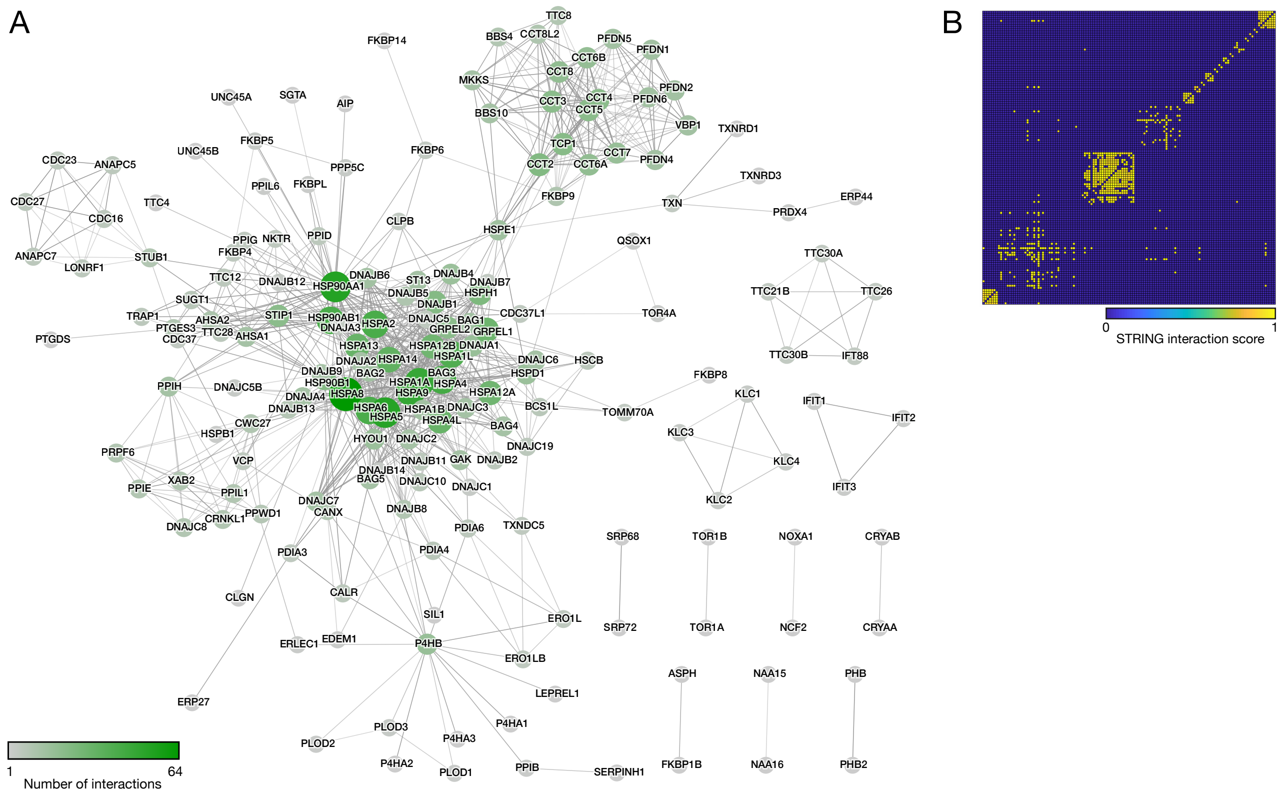


**Figure S2. Application of network analysis to define functional groups within the human chaperome.** (**A**) The 332 protein human chaperone network (Brehme et al, 2014), with nodes coloured according to chaperone degree as a measure of importance to the function of the network (Rubinov & Sporns, 2010). The most highly connected proteins in the network were: HSPA8 (degree = 64); HSP90AA1 (53); HSPA5 (52); HSPA9 (48); and, HSPA1A (47). (**B**) The adjacency matrix of the human chaperone network following modularity analysis. Rows and columns are the 332 nodes in the network, with entries representing the STRING interaction score (Szklarczyk et al, 2019). Interactions have been filtered to only include those of the highest confidence (score ≥ 0.9).

**Figure S3.**

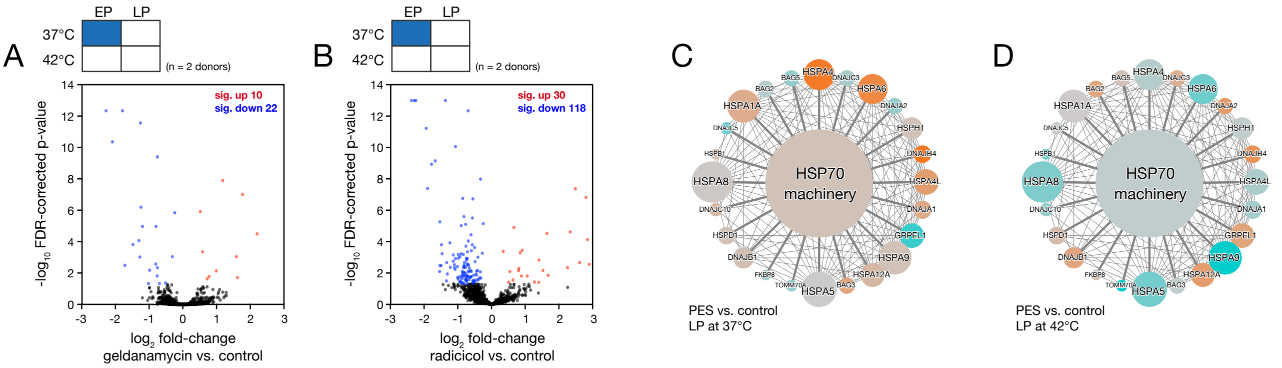


**Figure S3. Proteomic analysis of early passage (EP) and late passage (LP) primary human mesenchymal stem cells (hMSCs) treated with chaperone inhibitors and thermal stress.** Volcano plots showing changes to the proteomes of EP hMSCs treated with the HSP90-inhibitors (**A**) geldanamycin, or (**B**) radicicol, at 37 °C. Red and blue points satisfy a *p*-value < 0.05. *p*-values were calculated using empirical Bayes-modified t-tests with Benjamini–Hochberg false discovery rate (FDR) correction (Mallikarjun et al, 2020). (**C**) Network of proteins in the HSP70 chaperome module, showing changes induced by HSP70-inhibitor 2-phenylethynesulfonamide (PES) in LP hMSCs under control conditions. (D) Changes to protein levels in the HSP70 chaperome module induced by PES treatment in LP hMSCs subjected to 2-hour heat treatment at 42 °C. In network diagrams, node size is indicative of chaperone degree, while edge weight indicates interaction score between chaperones. The central node is coloured according to the mean abundance change of chaperones associated with the module. Descriptions of the chaperone modules can be found in Fig. 2A of the main paper.

**Figure S4.**

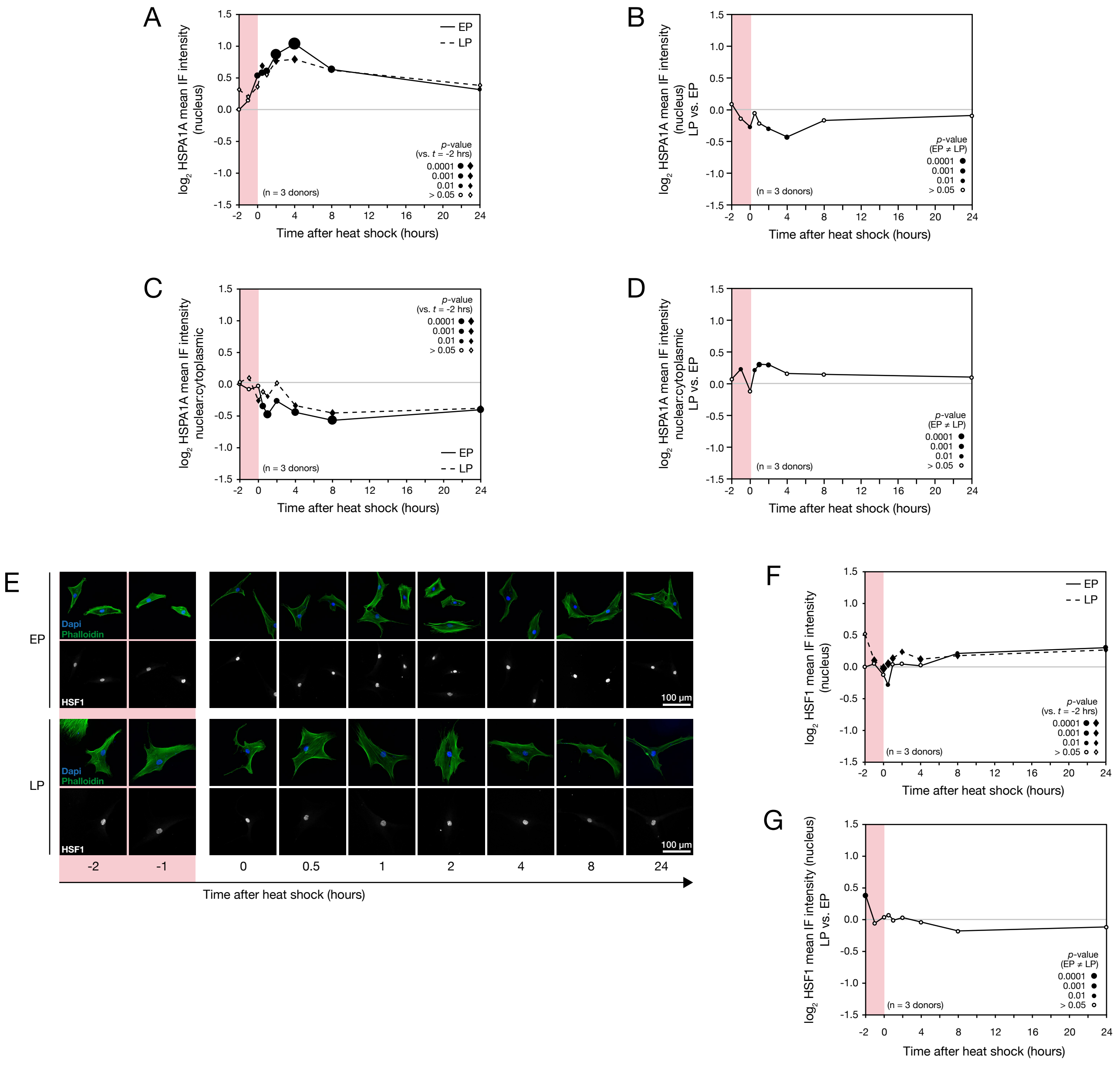
A
B
C
D
E
F
G
log2 HSPA1A mean IF intensity (nucleus)
log2 HSPA1A mean IF intensity (nucleus) LP vs. EP
log2 HSPA1A mean IF intensity nuclear:cytoplasmic
log2 HSPA1A mean IF intensity nuclear:cytoplasmic LP vs. EP
log2 HSF1 mean IF intensity (nucleus)
log2 HSF1 mean IF intensity (nucleus) LP vs. EP
Time after heat shock (hours)
EP
LP
p-value (vs. t = -2 hrs)
p-value (EP ≠ LP)
0.0001
0.001
0.01
> 0.05
(n = 3 donors)
Dapi
Phalloidin
HSF1
100 µm
-2
-1
0
0.5
1
2
4
8
24

**Figure S4. Analysis of immunofluorescence (IF) images of heat shock protein 70 kDa (HSPA1A) and heat shock factor 1 (HSF1) in early passage (EP) and late passage (LP) human mesenchymal stem cells (hMSCs) subjected to heat treatment.** hMSCs were imaged before, during and over a 24-hour period following a 2-hour heat treatment at 42 °C (see Fig. 4A in the main paper). Mean HSPA1A intensities were determined in nuclear and cytoplasmic cellular regions, defined by areas of DAPI and phalloidin staining, respectively. (**A**) Levels of HSPA1A protein in the nuclei of EP and LP hMSCs before, during and after heat stress. (**B**) Ratios of mean intensities of HSPA1A in the nuclei of LP vs. EP hMSCs subjected to heat stress. (**C**) Nuclear-to-cytoplasmic ratios of HSPA1A in EP and LP hMSCs subjected to heat stress. (**D**) Nuclear-to-cytoplasmic ratios of HSPA1A in LP vs. EP hMSCs subjected to heat stress. (**E**) Representative IF images of HSF1 in EP and LP hMSCs before, during and for 24 hours following a 2-hour heat treatment at 42 °C. (**F**) Quantification of levels of HSF1 protein in the nuclei of EP and LP hMSCs before, during and after heat stress. (**G**) Ratios of mean intensities of HSF1 in the nuclei of LP vs. EP hMSCs subjected to heat stress. In figure panels (A), (C) and (F), point size indicates the significance of changes vs. pre-heating, with solid points showing $p < 0.05$ (from ANOVA testing, $n = 3$ donors). In panels (B), (D) and (G), point size indicates significance where EP ≠ LP, with solid points showing $p < 0.05$ (from ANOVA testing, $n = 3$ donors).

### Figure S5.

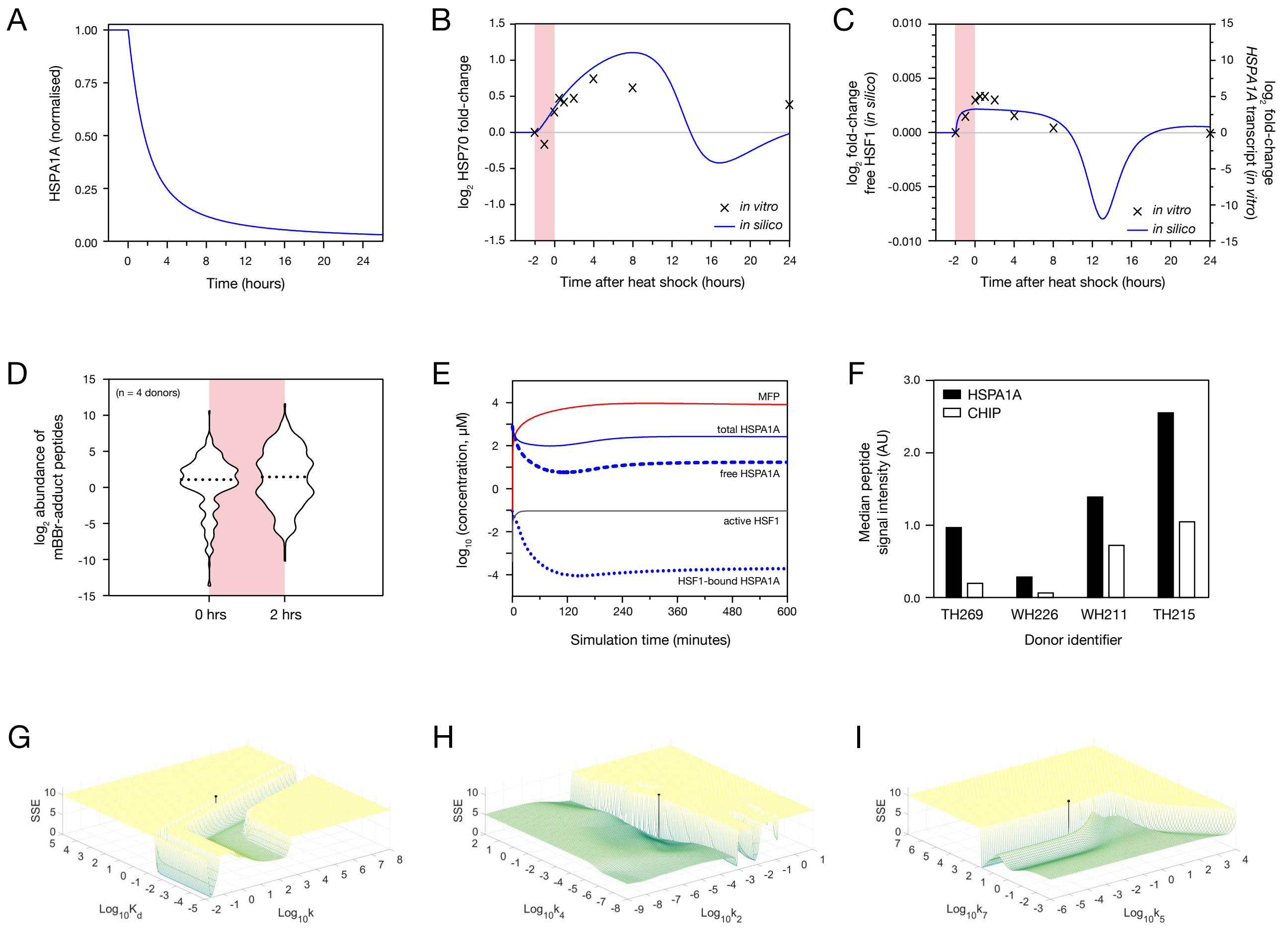

A
HSPA1A (normalised)
Time (hours)
B
log2 HSP70 fold-change
Time after heat shock (hours)
in vitro
in silico
C
log2 fold-change free HSF1 (in silico)
log2 fold-change HSPA1A transcript (in vitro)
Time after heat shock (hours)
in vitro
in silico
D
log2 abundance of mBBr-adduct peptides
(n = 4 donors)
0 hrs
2 hrs
E
log10 (concentration, μM)
MFP
total HSPA1A
free HSPA1A
active HSF1
HSF1-bound HSPA1A
Simulation time (minutes)
F
Median peptide signal intensity (AU)
HSPA1A
CHIP
TH269
WH226
WH211
TH215
Donor identifier
G
SSE
Log10Kd
Log10k
H
SSE
Log10k4
Log10k2
I
SSE
Log10k7
Log10k5

**Figure S5. Results supporting development and characterisation of an ordinary / delayed differential equation (ODE/DDE) model of the cellular response to heat stress.** (**A**) Decay of HSPA1A levels simulated by removing HSPA1A synthesis from the ODE/DDE model at equilibrium. The *in silico* half-life of HSPA1A was 100 minutes. (**B**) ODE/DDE model simulation of the stress response with adjustments made to resemble senescent cells. The *in silico* model was given sufficient time to reach a stable equilibrium, before a proteotoxic stress was simulated by increasing the rate at which misfolded proteins were generated within the model for 120 minutes. At each time interval, the *in silico* concentration of HSPA1A was recorded, and is shown overlaid with data acquired by immunofluorescence from late passage (LP) hMSCs *in vitro* (see Fig. 4B of the main paper). (**C**) As (B), but showing concentrations of modelled active HSF1 overlaid, with a scaling factor, onto experimentally-derived *HSPA1A* transcript levels in early passage (EP) hMSCs (see Fig. 4D of the main paper). (**D**) Abundances of monobromobimane (mBBr)-tagged peptides in mass spectrometry data pre-stress and immediately following a 2-hour treatment at 42 °C. Dotted lines indicate sample medians. (**E**) Convergence of populations in the ODE/DDE model to a stable equilibrium prior to stress. (**F**) HSPA1A and CHIP protein signal intensities quantified by mass spectrometry, in the absence of stress, in EP hMSCs from four human donors. (**G-I**) Demonstration of the optimisation of free model parameters. The ODE/DDE model was used to simulate the stress response over a range of parameter values, and the sum of squared errors (SSE) compared to experimentally derived data (measurements of HSPA1A by immunofluorescence) at each parameter value. The parameter values chosen by optimisation are shown overlaid in black over the SSE values.

**SUPPLEMENTAL TABLE**

| Parameter | Description | Value | Unit |
|---|---|---|---|
| $k$ | Maximal HSPA1A transcription rate | $6.4294 \times 10^{2}$ | $min^{-1}$ |
| $k$ | (Modified value in senescent cells) | $4.5865 \times 10^{2}$ | $min^{-1}$ |
| $n$ | Hill coefficient | $3$ | (dimensionless) |
| $K_d$ | HSF1 dissociation constant | $3.0534$ | $\mu M$ |
| $k_2$ | HSF1-HSPA1A binding rate | $3.9827 \times 10^{-4}$ | $\mu M^{-1} min^{-1}$ |
| $k_3$ | HSPA1A-MFP binding rate | $3.9827 \times 10^{-4}$ | $\mu M^{-1} min^{-1}$ |
| $k_4$ | CHIP-mediated HSPA1A turnover rate | $1.2400 \times 10^{-3}$ | $\mu M^{-1} min^{-1}$ |
| $k_5$ | HSPA1A-mediated MFP refolding rate | $5.0199 \times 10^{-2}$ | $min^{-1}$ |
| $k_6$ | CHIP-mediated MFP turnover rate | $2.1200 \times 10^{-3}$ | $\mu M^{-1} min^{-1}$ |
| $k_7$ | Rate of protein misfolding at 37 °C | $69.1068$ | $\mu M\ min^{-1}$ |
| $\lambda$ | Rate of protein misfolding at 42 °C | $4.2651 \times 10^{2}$ | $\mu M\ min^{-1}$ |
| $t$ | Simulation time elapsed | | $min$ |
| $\tau$ | HSF1 activation/HSPA1A synthesis delay | $\sim U[60, 180]$ | $min$ |

**Supplemental Table S6.** Optimised and approximated parameters for the ordinary/delay differential equation (ODE/DDE) mathematical model of the heat stress response. MFP = misfolded protein; CHIP = E3 ubiquitin-protein ligase CHIP.

### SUPPLEMENTAL REFERENCES

Brehme M, Voisine C, Rolland T, Wachi S, Soper JH, Zhu Y, Orton K, Villella A, Garza D, Vidal M, Ge H, Morimoto RI (2014) A chaperome subnetwork safeguards proteostasis in aging and neurodegenerative disease. *Cell Reports* **9:** 1135-1150

Brocchieri L, Karlin S (2005) Protein length in eukaryotic and prokaryotic proteomes. *Nucleic Acids Research* **33:** 3390-3400

Hernandez-Segura A, Nehme J, Demaria M (2018) Hallmarks of cellular senescence. *Trends in Cell Biology* **28:** 436-453

Mallikarjun V, Richardson SM, Swift J (2020) BayesENproteomics: Bayesian elastic nets for quantification of peptidoforms in complex samples. *Journal of Proteome Research* **19:** 2167-2184

Rubinov M, Sporns O (2010) Complex network measures of brain connectivity: Uses and interpretations. *Neuroimage* **52:** 1059-1069

Shimi T, Butin-Israeli V, Adam SA, Hamanaka RB, Goldman AE, Lucas CA, Shumaker DK, Kosak ST, Chandel NS, Goldman RD (2011) The role of nuclear lamin B1 in cell proliferation and senescence. *Genes & Development* **25:** 2579-2593

Szklarczyk D, Gable AL, Lyon D, Junge A, Wyder S, Huerta-Cepas J, Simonovic M, Doncheva NT, Morris JH, Bork P, Jensen LJ, Mering C (2019) STRING v11: protein-protein association networks with increased coverage, supporting functional discovery in genome-wide experimental datasets. *Nucleic Acids Research* **47:** D607-D613
